# Evolutionary Consequences of Unusually Large Pericentric TE-rich Regions in the Genome of a Neotropical Fig Wasp

**DOI:** 10.1101/2025.04.20.649723

**Authors:** Zexuan Zhao, Kevin Quinteros, Carlos A. Machado

## Abstract

Transposable elements (TEs), despite generally being considered deleterious, represent a substantial portion of most eukaryotic genomes. Specific genomic regions, such as telomeres and pericentromeres, are often densely populated with TEs. In these regions, which tend to be gene-poor, reduced recombination shelters the genome from the deleterious effects of TEs. Here, we describe unusually large and continuous pericentromeric TE-rich regions in all chromosomes of the genome assembly of *Pegoscapus hoffmeyeri* Sp. A (511.79 Mbp), a Neotropical fig wasp that is the obligate pollinator of *Ficus obtusifolia*. The identified pericentromeric TE-rich regions span nearly half (46%) of the genome, and harbor over 40% of all annotated genes, including 30% of conserved BUSCO genes. We present evidence that low recombination in these TE-rich regions generates strong bimodal molecular evolution patterns genome-wide. Patterns of nucleotide diversity and protein-coding gene evolution in TE-rich regions are consistent with a reduced efficiency of selection and suggestive of strong Hill–Robertson effects. A significant reduction in third codon position GC content (GC3) in TE-rich regions emerged as the most distinctive gene feature differentiating genes in TE-rich regions from those in the rest of the genome, a pattern that likely results from the absence of GC-biased gene conversion. This remarkable binary compartmental genome organization in the genome of *P*. *hoffmeyeri* provides a unique example of how genome organization with compartmental TE distribution can lead to context-dependent gene evolution shaped by common evolutionary forces.

## Introduction

Haploid genome sizes of eukaryote species show little correlation with organismal complexity (Lynch and Conery 2003; Wright 2017; Blommaert 2020). In insects, haploid genome sizes can vary by two orders of magnitude, ranging from 89.6 Mbp in the Antarctic midge, *Belgica antarctica* (Kelley et al. 2014) to 21.48 Gbp in the grasshopper *Bryodemella tuberculata* (Hawlitschek et al. 2023). Given that the number of genes remain relatively conserved across taxa, discrepancies in haploid genome sizes are predominantly the result of differences in the extent and composition of intronic and intergenic sequences (Lynch and Conery 2003; Elliott and Gregory 2015a). As transposable elements (TEs) are major components within these regions (Bourque et al. 2018), they have become a major focus of research into the mechanisms underlying the evolution of genome structure (Chalopin et al. 2015; Elliott and Gregory 2015b; Sessegolo et al. 2016; Kapusta et al. 2017; Blommaert et al. 2019; Cong et al. 2022; Zuo et al. 2023; Betancourt et al. 2024).

Although some studies have reported beneficial effects of TEs through their effects on regulatory evolution, chromosome structure, or horizontal gene transfer (Klein and O’Neill 2018; Choudhary et al. 2020; Widen et al. 2023), most TEs are thought to be deleterious and to be under purifying selection (Cridland et al. 2013; Blumenstiel et al. 2014; Stritt et al. 2018; Oggenfuss et al. 2021; Dazenière et al. 2022; Betancourt et al. 2024). Newly inserted TEs can disrupt coding sequences (Charlesworth and Langley 1989; Hancks and Kazazian 2016), and remnant TE copies can trigger ectopic recombination and chromosomal rearrangement between non-homologous sequences across chromosomes (Montgomery et al. 1987; Langley et al. 1988), dramatically increasing the risk of deleterious effects as TE copy number increases (Bennetzen and Wang 2014). Even when TEs are suppressed through epigenetic modification, the inadvertent spreading of these modifications can negatively affect the expression of neighboring genes (Choi and Lee 2020). Despite these potentially harmful effects associated with TEs, their prevalence across genomes remains a puzzling paradox that underscores their complex evolutionary and functional significance.

One explanation for the prevalence of TEs is the existence of “TE refuges” in genomes (Bertocchi et al. 2018). For instance, the Mbp-scale pericentric heterochromatic regions flanking centromeres in *Drosophila melanogaster* are populated with satellite DNA and TEs and even contain genes of exceptionally large size, such as *Myo81F*, which spans over 2.5 Mbp (Hoskins et al. 2007; Hoskins et al. 2015). Other examples include non-recombining regions of sex chromosomes (Erlandsson et al. 2000; Duhamel et al. 2023), B chromosomes (Bertocchi et al. 2018), heterochromatic knobs (Quesneville 2020) and chromosomal inversions (Carpinteyro-Ponce and Machado 2024). These regions, characterized by suppressed recombination, limit the deleterious effects of ectopic recombination and/or slow down the removal of TEs through recombination (Kent et al. 2017). Consequently, these genomic environments often harbor TE families with higher copy numbers, while other gene-rich regions accumulate younger TE families (Baucom et al. 2009). While TEs can accumulate in regions with low recombination, TE silencing could also further reduce recombination rates by introducing repressive chromatin modifications (Mirouze et al. 2012; Yelina et al. 2012; Zamudio et al. 2015; Huang et al. 2025). In addition, the coevolution between TEs, recombination and chromatin structure can also influence gene evolution in direct or indirect ways. TEs can directly influence gene evolution in TE rich regions through their accumulation in introns, resulting in increased gene size (Yasuhara et al. 2005; Corradini et al. 2007). Further, in regions of low recombination the efficacy of selection is weakened due to increased Hill–Robertson interference (Hill and Robertson 1966), leading to higher sequence divergence of protein coding genes in TE-rich regions (Campos et al. 2012). The intricate interplay between TEs and chromatin structure, as well as their compound effects on heterochromatic genes, need to be further investigated.

In this study, we present a comprehensive analysis of the evolutionary effects of TEs in the genome of one pollinating fig wasp (Superfamily Chalcidoidea, family Agaonidae). The karyotype of Agaonids is relatively stable at 5-6 chromosomes across the family, and most chromosomes are metacentric (Liu et al. 2011; Gokhman et al. 2019). The conserved chromosome structure and lack of large-scale chromosome fusions or translocations provide a structurally stable genomic landscape to detect and track the evolution of TE refuges. Fig pollinating wasps are highly specialized organisms that spend most of their life cycle (3-4 weeks) developing in fig tissue, and only 1-2 days as adults searching for a new receptive fig host (Janzen 1979; Weiblen 2002). Based on their special ecological niche, fig wasps were used to support the hypothesis that TEs should experience limited accumulation in genomes of specialists whose ecological contacts with other organisms are rare (Gilbert et al. 2021). In fig wasps, this hypothesis was initially supported by the short-read based genome assembly of *Ceratosolen solmsi* which showed very low (6.4%) TE content (Xiao et al. 2013). However, as genome assemblies for fig wasps have become more refined, the estimated TE content has approached 20% (Cooper et al. 2020; Zhang et al. 2020), challenging this original hypothesis. In this study, we further challenge this idea by generating a chromosome-level genome assembly of *Pegoscapus hoffmeyeri* sp. A (Grandi 1934), one of the two pollinator species associated with *Ficus obtusifolia* in Panama (Molbo et al. 2003; Molbo et al. 2004). The assembled genome is 30-85% larger than published genomes from other fig wasp species due a remarkable increase in TE content. We show that TE content in this species is effectively bimodal, with extremely large pericentric TE-rich regions encompassing almost half of the genome and protein coding genes, and showing patterns of molecular evolution markedly differently than the rest of the genome.

## Materials and Methods

### High Molecular DNA Extraction and Reference Genome Sequencing

We collected a brood from a fig pollinated by a single foundress on Barro Colorado Island, managed by the Smithsonian Tropical Research Institute. High molecular weight DNA was extracted using the Qiagen Blood and Cell culture Midi Kit following a previously described protocol (Chakraborty et al. 2016). The Pacbio HiFi library was constructed and sequenced in a Sequel II instrument at the Institute for Genome Sciences (University of Maryland).

### Genome Assembly and Quality Assessment

The raw subreads were aligned using CCS algorithm v 6.4.0 (https://github.com/PacificBiosciences/ccs) to call high-fidelity (HiFi) long reads. Contigs were assembled by Hifiasm v0.15 (Cheng et al. 2021) and the taxonomical match of each contig was identified through Basic Local Alignment Search Tool (BLAST) against the nt database (Altschul et al. 1990; Sayers et al. 2022). Contigs taxonomized as insects or unclassified were scaffolded using RagTag v1.0.2 (Alonge et al. 2022) using the genome assembly of *Eupristina verticillata* (Accession # GWHFQFG00000000.1) as a reference. Assembly duplication was checked by self-alignment dot plots using D-Genies v1.5.0 (Cabanettes and Klopp 2018). HiFi reads were mapped to the assembly using minimap2 v2.26 (Li 2018; Li 2021) and mapping depths were calculated using samtools v1.17 (Li et al. 2009; Danecek et al. 2021).

The quality of the genome assembly was evaluated using the Genome Evaluation Pipeline (https://git.imp.fu-berlin.de/cmazzoni/GEP). It used the Python script ‘assembly_stats.py’ to calculate scaffold and contig statistics from the assembled genome (Trizna 2020). GenomeScope v2.0 estimated genome size, repeat content, and heterozygosity from HiFi reads (Vurture et al. 2017; Ranallo-Benavidez et al. 2020). Merqury v1.3 (Rhie et al. 2020) evaluated kmer completeness, accuracy, and heterozygosity of the assembly, while BUSCO v5.4.7 (Manni et al. 2021; Alonge et al. 2022) assessed gene completeness against the hymenoptera_odb10.2019-11-20 database. To examine whether the assembly reached pseudochromosome level, telomeric repeats were identified using tidk v0.2.3 (Brown et al. 2023).

### Genome Annotation

De novo transposable element annotation was performed using Extensive de novo TE Annotator (EDTA) v2.1.0 (Ou et al. 2019). EDTA incorporates a variety of repeat annotators into its pipeline: GenomeTools v1.6.2 (Gremme et al. 2013), LTR_FINDER v1.07 (Xu and Wang 2007), LTR_retriever v2.9.0 (Ou and Jiang 2018), Generic Repeat Finder v1.0 (Shi and Liang 2019), TIR-Learner v2.5 (Su et al. 2019), HelitronScanner v1.1 (Xiong et al. 2014) and TEsorter v1.3 (Zhang et al. 2022). It also uses RepeatModeler v2.0.3 (Flynn et al. 2020) to identify remaining TEs and masks the genome using RepeatMasker v4.1.2 (http://www.repeatmasker.org). For comparison, we also annotated the genome assemblies of *V. javana*, *W. pumilae*, *C. solmsi marchali* and *E. verticillata* to control the different performance of TE annotation pipelines. We classified TEs into superfamilies according to (Wicker et al. 2007), and assigned TEs that were classified only into order or higher classification level or unclassified as unclassified TEs. The *createRepeatLandscape.pl* script from RepeatMasker was used to generate a TE landscape.

We employed homology-based and de novo gene prediction on the genome assembly with only long TEs (length >=1 Kb) masked. Homology-based gene prediction was done using Gene Model Mapper (GeMoMA) v1.9 (Keilwagen et al. 2019) with 10 Hymenoptera genome annotations as references with a variety of genetic distances (Supplementary table 5). We turned off static intron length (*GeMoMa.sil=false*) in GeneModelMapper step to infer intron length from reference annotation individually for each gene. It is worth noting that GeMoMa tends to overestimate genes when multiple references are provided. To eliminate bias towards genes with extremely long sequences and control false discovery rate, we tested *score/aa* ratio from 0.8 to 2.0 with increment = 0.1 in GeMoMa’s Annotation Filter step and evaluated the quality of annotation using BUSCO completeness. We picked score/aa=1.5 as it generated the highest BUSCO completeness (Supplementary table 6). Using this optimized filtering parameter we predicted 12,285 genes, compared to 16,332 genes predicted using the default filter. De novo gene prediction was done using Augustus v3.5.0 (Stanke et al. 2006). We used 1000 genes predicted by GeMoMa as a validation set and the rest as the training set to train and validate a de novo gene model. In total 23,777 genes were predicted by Augustus, of which 13,208 genes do not overlap with GeMoMa predicted genes. We used eggNOG-mapper v2.1.9 (Cantalapiedra et al. 2021) with eggNOG database v5.0 (Huerta-Cepas et al. 2019) for functional annotation of the predicted genes. A total of 10,264 genes from the 12,285 GeMoMA predicted genes and 2,872 from the 13,208 de novo predicted genes were functionally annotated. In total, we annotated 13,136 functional genes using homology-based and de novo approaches. Based on the longest isoforms for each genes, we also annotated the non-degenerate and four-fold degenerate sites using degenotate v1.3 (Mirchandani et al. 2024).

### Classification of TE-rich Regions

We calculated TE densities as the proportion of base pairs covered by homology-based TE annotations in 200-Kbp genomic windows after merging overlapping TE annotations using the GenomicRanges package v3.17 (Lawrence et al. 2013) in R v4.3.0 (R Core Team 2023). As the frequency of TE densities suggests a bimodal distribution (Supplementary figure 8), we fitted mixture distributions of k subpopulations with k from 1 to 3 to the TE densities, and calculated Akaike information criterion (AIC) and Bayesian Information Criterion (BIC) values to select the best mixture model.

One assumption to classify genomic windows is local correlation. If the TE density of a window falls between two mean TE density peaks, the classification should rely on the classification of neighboring regions. To employ this idea, we fitted the TE densities along the five chromosomes to a hidden Markov model (HMM) with two states using depmixS4 v1.5.0 (Visser and Speekenbrink 2010) package in R. HMM utilizes both the magnitude of TE densities and the transition of the densities to infer the state of each window. Two states with different mean TE densities can switch along the genome according to a transition probability matrix. The inter-state transition probability represents the probability of switching to a TE-rich window from a background window or vice versa. The lower the transition probability, the bigger the expected TE-rich islands, and a higher level of clustering pattern. We estimated the significance of the clustering pattern by permuting TE densities on the chromosomes and fitting the HMM to the permuted datasets. The p-value of observed transition probability is calculated from the distribution of permuted transition probabilities.

### Whole Genome Resequencing and Estimation of Nucleotide Diversity

We collected eight female offspring from independent fig syconia from Panama’s Barro Colorado Monument. DNA was extracted using Omega Biotek’s E.Z.N.A. Mollusc & Insect DNA Kit. Illumina libraries were prepared using the NEBNext Ultra II FS DNA Library Prep Kit for Illumina (New England Biolabs) and libraries were sequenced on an Illumina HiSeq 3000 instrument at the Institute for Genome Sciences (University of Maryland).

We compiled a customized workflow to detect SNPs from the Illumina short read population genomic dataset. Low-quality bases and adaptor sequences were trimmed by fastp v0.23.2 (Chen et al. 2018; Chen 2023). The quality of reads was assessed using fastQC v0.23.2 (Andrews et al. 2012) before and after trimming. Using the ‘fq2bam’ command within clara-parabricks v4.0.0 (https://docs.nvidia.com/clara/parabricks/4.0.0/index.html), each sample was aligned to the reference genome and duplicates were marked. Average read depth was estimated as the number of reads overlapping each base pair, and average read coverage was estimated as the proportion of base pairs covered by at least one read are examined across the genome and samples (Supplementary figure 9).

Nucleotide diversity (π) is the average pairwise difference between all possible pairs of individuals (Nei and Li 1979) and is normalized by the length of the sequences being compared. In the traditional variant calling pipeline, missing data is treated as reference calls, and when calculating π, the denominator used is the length of the sliding windows rather than the actual length of sequences observed, which can potentially lead to an underestimation of π. To correct the bias, we called genotypes at all sites in the reference genome. Variants were generated by DeepVariant 1.4.0-gpu (Poplin et al. 2018), and jointly genotyped by GLnexus v1.4.1 (Yun et al. 2021), and biallelic SNPs were selected by BCFtools v1.17 (Danecek et al. 2021). We used DeepVariant for its high accuracy in genotype calling, while it does not support calling reference loci. Therefore, we generated reference genotype calls by BCFtools v1.17, and merged variant calls and reference calls using bedtools v2.31.0 (Quinlan and Hall 2010) and GATK4 v4.4.0.0 (Quinlan and Hall 2010). We filtered variant and reference genotype calls by vcftools v0.1.16 (Danecek et al. 2011) with parameter *--max-missing 0.8 --min-meanDP 15 --max-meanDP 50 -- minDP 10 --maxDP 100*. We further separated genotype calls by gene annotation and TE annotation using bedtools. Nucleotide diversity was calculated using pixy v1.2.7.beta1 (Korunes and Samuk 2021) per 200-Kbp non-overlapping sliding windows. We added the total number of reference calls and variant calls of respective categories in each window as a confounding variable. For comparison, we also calculated uncorrected π from variant calls straight from DeepVariant using vcftools.

To assess whether our sequencing depths are sufficient for accurate genotyping, we down-sampled reads to 10x and 20x coverage using SeqKit v2.5.1 (Shen et al. 2016) and calculated recall, precision and F1 scores against variant calls derived from the full set of reads using Haplotype Comparison Tools v0.3.15 (https://github.com/Illumina/hap.py).

### CpG Methylation Detection

5-Methylcytosine (5mC) at CpG sites was first detected on the PacBio HiFi reads from subreads using jasmine v2.0.0 (https://github.com/pacificbiosciences/jasmine). Then these reads were aligned to the genome using pbmm2 v1.13.1 (https://github.com/PacificBiosciences/pbmm2). For each CpG in the reference genome, the per-site methylation probability was calculated using pb-CpG-tools v2.3.2 (https://github.com/PacificBiosciences/pb-CpG-tools). To establish an optimal probability threshold to classify methylated CpGs, we plotted candidate probability thresholds against the detected number of CpG methylation sites. The optimal threshold (0.65) was identified by locating the threshold value where the absolute value of its local slope is at its minimum (Supplementary figure 13). We assigned CpGs to their nearest genes or TEs to avoid redundancy, and then counted the number of methylated CpGs in 500-bp windows. We then performed principal component analysis (PCA) and determined windows with the highest 5% loadings of PC1 as regions with differentially methylated CpGs. To validate the CpG methylation detection, we tested if lower observed-to-expected CpG ratios of methylated genes were observed than expected (Supplementary figure 16). Observed-to-expected CpG ratios in coding sequences were calculated using the formula from (Elango et al. 2009).

### Summary of Gene Features

The following additional features were tabulated as the input for the machine learning model: number of exons, number of introns, intron GC content, third codon position GC content (GC3), first and second codon position GC content (GC12), coding sequence lengths, total intron lengths, and number of TE-insertions in introns. We also included maximum likelihood codon bias (MCB) estimated using coRdon package v1.20.0 in R (Elek et al. 2024). Synonymous divergence (dS), non-synonymous divergence (dN) and non-synonymous to synonymous divergence ratio (dN/dS) was calculated against protein sequences from another fig wasp *E. verticillata* using orthologr package v0.4.2 in R (Drost et al. 2015), in which reciprocal BLAST hits were used to detect orthologs and the Comeron method was used to estimate dN and dS (Comeron 1995). We also included gene annotation method (source) as a gene feature to avoid bias in annotation accuracy.

### Random Forest Analysis and Importance Analysis of Gene Features

Random forest, a naive machine learning algorithm, implemented in randomForest R package v4.7-1.1 (Liaw and Wiener 2002) was used to predict whether genes were in TE-rich regions or other regions from aforementioned gene features, and to determine which features were most important. It uses an ensemble of decision trees to predict the response, each of which was built upon a random selection of features. To avoid overfitting, we only trained shallow trees (maximum number of terminal nodes = 5). We randomly selected 80% of genes to train the model and used the rest to test its performance. To avoid bias, the dataset was resampled to balance the number of genes in the TE-rich regions and other regions. The importance of gene features was measured as increased error rate after permuting each feature.

## Results

### *Pegoscapus hoffmeyeri* sp. A has an unexpectedly large genome with high TE content

From an extremely rare all-male brood from a single *P. hoffmeyeri* sp. A foundress, we generated 11.9 Gbp from 1.34M PacBio HiFi reads, expecting 40x coverage according to the sizes of genome assemblies of other fig wasps (around 300 Mbp) (Table 1). However, k-mer analysis (k=31) of the HiFi reads suggests that the haploid genome size is 500.96 Mbp or 40% larger than other published fig wasp genomes, with extremely low heterozygosity (0.001%) (Supplementary figure 1). The low heterozygosity observed in *P. hoffmeyeri* sp. A is consistent with the highly inbred nature of *P. hoffmeyeri* sp. A (Molbo et al. 2004). However, the large genome size was unexpected and was further investigated to determine if it was overestimated.

**Table 1:**
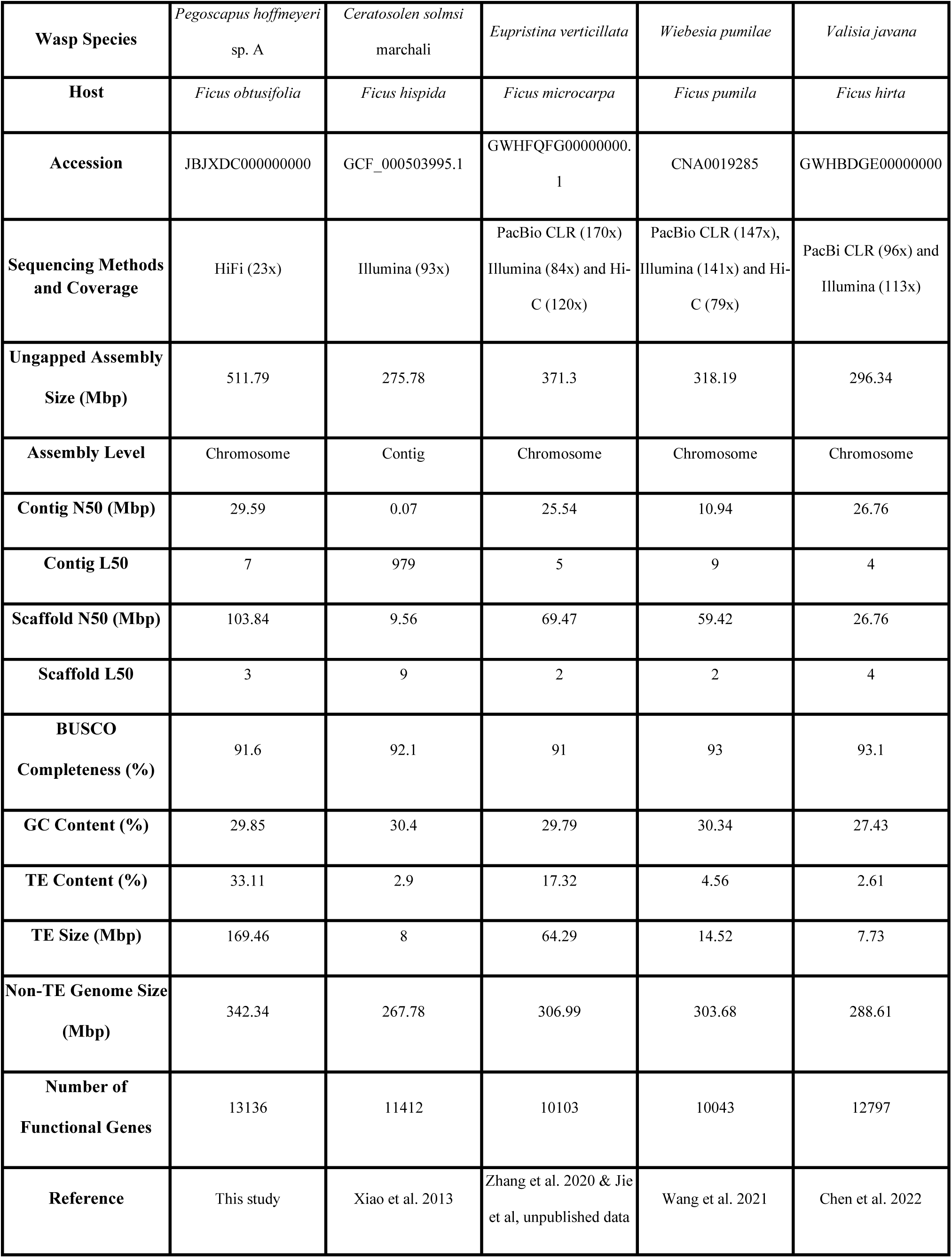
Statistics of the *P. hoffmeyeri sp.A* genome assembly, and comparison with other fig wasp genome assemblies.

The initial contig-level assembly had a length of 511.79 Mbp with a contig N50 of 29.59 Mbp. These findings were consistent with our assembly-free k-mer estimations. Using an assembly from *Eupristina verticillata* as a reference (Jie et al, unpublished data), we scaffolded 65 contigs onto 5 chromosomes with a scaffold N50 of 103.83 Mbp while only 1.5% of the initial assembly remained unplaced (Supplementary figure 2A, Supplementary figure 3, Table 1). Self-alignment of the genome assembly showed no significant duplicated segments in the assembly (Supplementary figure 4A). Benchmarking Universal Single-Copy Orthologs (BUSCO) analysis showed that the assembly contained 5489 out of 5991 (91.6%) complete Hymenoptera orthologs, with only 36 identified (0.6%) duplicates. The BUSCO completeness is consistent with those observed in other fig wasp genome assemblies (Table 1). K-mer analysis indicated that 99.89% of 31-mers present in the HiFi reads are in the assembly, indicating a low level of misassembly such as switch errors. Mapping depths were consistent across the genome (Supplementary figure 4B), except for some sparse regions on the 5’ end of chromosome 1. Altogether these analyses suggest that the estimation of genome size was not inflated and thus the genome of *P. hoffmeyeri* sp. A is 30-85% larger than previously published fig wasp genomes (Table 1).

We conducted further assessment of assembly quality through whole genome annotation. Utilizing a homology-based annotation method (GeMoMA) (Keilwagen et al. 2019), we identified 12,285 genes using a database compiled from mosquitoes, bees, flies, and other fig wasps (Supplementary table 5) (Friedrich and Muqim 2003; Xiao et al. 2013; Hoskins et al. 2015; Wallberg et al. 2019; Dalla Benetta et al. 2020; Zhang et al. 2020; Wang et al. 2021; Habtewold et al. 2023; Toga et al. 2024; Lee et al. 2025). In addition, we predicted 23,777 genes using a *de novo* approach. We removed *de novo* predicted genes that overlapped with homology-based genes and annotated the function of the remaining set of genes. The final gene set comprises 13,136 functionally annotated protein-coding genes, which is comparable to other fig wasp genomes (Supplementary figure 2b, Table 1). Additionally, we annotated 89 complete rRNA loci (29 5S, 19 5.8S, 20 18S and 21 28S), and 166 tRNA loci (Supplementary figure 2b). We also identified AACCCAAT/ATTGGGTT telomeric repeats at both ends on 4 chromosomal scaffolds (Supplementary figure 2b).

We annotated transposable elements (TEs) to investigate their contribution to the observed genome size expansion in *Pegoscapus hoffmeyeri* sp. A. We found that 33.11% of the genome is occupied by TEs (Figure 1A), the highest percentage among all sequenced fig wasp genomes to date. The genome size after excluding TEs is 340 Mbp. This number aligns more closely with genome sizes after TE removal that have been reported in other fig wasp species, which range from 267.78 to 317.79 Mbp (Table 1). The five most abundant classified TE superfamilies—Helitron, Gypsy, Mutator, Copia, and Tc1_Mariner—constitute 6.22%, 5.33%, 3.31%, 2.29%, and 2.29% of the genome, respectively. When compared to *E. verticillata* (Supplementary figure 5A), *P. hoffmeyeri* sp. A possesses a larger quantity of DNA transposons, with a notable increase in the Helitron superfamily.

**Figure 1.**
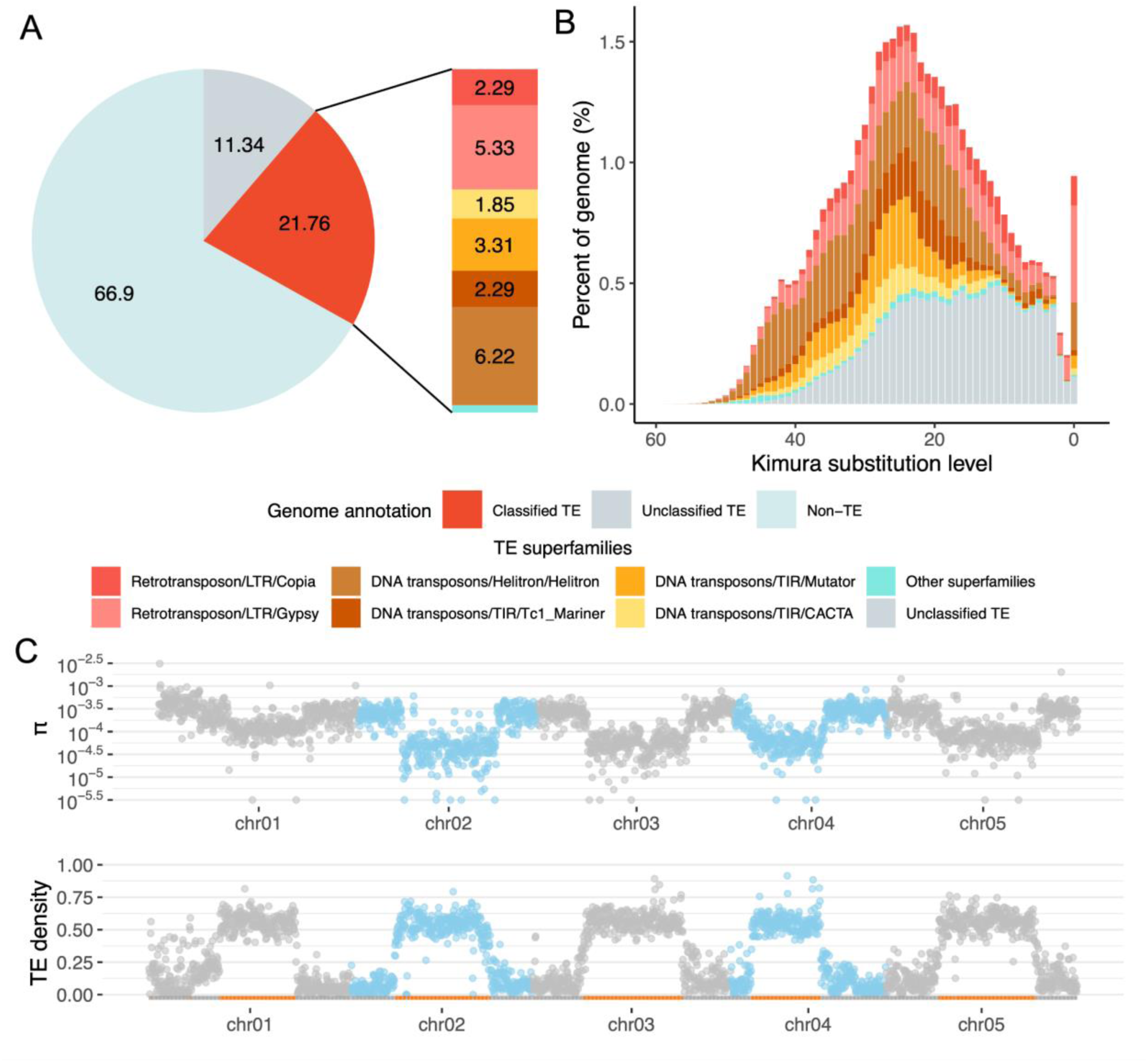
TE composition, landscape and distribution associated with nucleotide diversity. (A) Genome composition of TE and non-TE elements. (B) The TE landscape; the X-axis represents the pairwise CpG-corrected Kimura 2-parameter substitution level of the TEs relative to the consensus sequence of their respective families, analogous to the age of the TEs, while the Y-axis is the percentage of the genome occupied by TEs. (C) Patterns of nucleotide diversity and TE density, with classified TE-rich regions in orange on the X-axis.

To estimate the timeline of TE accumulation, we plotted the TE landscape, displaying the distribution of Kimura distances across TE superfamilies. A bimodal distribution was observed (Figure 1B), with a majority of TEs in the older part of the bell-shaped distribution. Using a mutation rate of *μ* = 6.8 × 10^−9^per site per generation estimated from honeybees (Yang et al. 2015) and 6 generations per year (Weiblen 2002), the expected Kimura substitution level is 4.08 per million years. The peak of the bell-shaped distribution is estimated to have occurred approximately 5.6 mya. There is another distinct peak near zero on the TE landscape, which suggests an ongoing transposition activity. The genome of *E. verticillata* shows a consistently steady and moderate TE activity and also an ongoing transposition activity on the TE landscape (Supplementary figure 5), but both processes have been or are less intense than those observed in *P. hoffmeyeri* sp. A.

One important consequence of the observed TE accumulation is its effect on gene size, specifically intron size. While only 17.54 Mbp of the genome are exons, 170.27 Mbp are introns (Supplementary table 1). Within these introns, 54.91 Mbp are classified as TEs, with 12.18 Mbp specifically identified as Helitrons. Genes with intronic TE insertions (median size = 10,825 bp, n = 6058) are an order of magnitude larger than genes without TE insertions (median size = 1,653 bp, n = 7078) (p<0.001, Wilcoxon test) (Supplementary figure 6).

### A Binary Genome-wide Pattern of TE Density, Nucleotide Diversity and Protein Coding Gene Evolution

To understand the organization of TEs across the genome, we calculated TE densities (see Methods) and explored their spatial pattern (Figure 1C). A binary pattern of TE densities was observed, and the rank orders of TE densities per superfamily were similar across the genome (Supplementary figure 7). A modality test supported a bimodal distribution (Supplementary figure 8, Supplementary table 2). A 2-state hidden Markov model trained solely by TE densities identified five major regions with elevated TE densities, amounting to 46.3 % of the genome (237 Mbp) (Figure 1C). Around 80% of TEs were located in these TE-rich regions. The average TE density is much higher in TE-rich regions (mean = 56.54%) compared to the other regions (mean = 11.60%). For each chromosome, TE-rich regions comprise 37.6 Mbp to 54.6 Mbp, or 38.56% to 51.02% of the chromosomes (Supplementary table 3). These regions are significantly clustered (p<0.001, permutation test) and major TE-rich regions are located close to the putative center of the chromosomes. Remarkably, these TE-rich regions also encompass 40.8% of all annotated genes, and are thus not gene-poor regions. In fact, the average gene density is only slightly decreased in TE-rich regions (4.26 genes per 200 Kbp) compared to other regions (5.53 genes per 200 Kbp) (p=0.0059, Wilcoxon test) (Supplementary figure 6).

To understand the effect of these TE-rich regions on molecular evolutionary processes, we first estimated nucleotide diversity (π) using whole genome short read sequencing from 8 individuals and compared π within and outside these TE-rich regions. 83.2M to 118.2M Illumina reads per individual were generated from 8 female wasps, providing 21x to 30x coverage of the genome (Supplementary table 4, Supplementary figure 9). In total, we called 239,435 SNPs using DeepVariant. Down-sampling analysis shows consistent genotype calls even when coverage was reduced to 10x, and thus the sequencing depth was sufficient for reliable SNP calling (Supplementary figure 10). In repetitive regions, short-read alignment is more challenging, resulting in a higher false negative rate in variant calling. To address this potential source of underestimation of π, we called 361.51M reference genotypes, and calculated π exclusively from these reference and variant positions with similar quality (Supplementary figure 11).

Similar to TE densities, nucleotide diversity (π) showed a binary pattern across the genome (Figures 1C). π is significantly lower in TE-rich regions (p < 2 x 10^-16^, Wilcoxon test), with a median π of 7.08 × 10^−5^, compared to a median π of 2.84 × 10^−4^in other regions. This pattern is also consistent across exons, intronic TEs, TE-free introns, intergenic TEs and intergenic non-TE sequences (Figure 2). We found that effective sequence size, the total number of reference and variant sites when calculating π, heavily influences π estimation. In intronic and intergenic regions, π stabilized when effective sequence size was above 20 Kbp and was consistently lower in TE-rich regions. Although in exonic regions π hardly stabilized because exons are too sparse, exons in TE-rich regions still exhibited the lowest level of π compared to other sequences.

**Figure 2.**
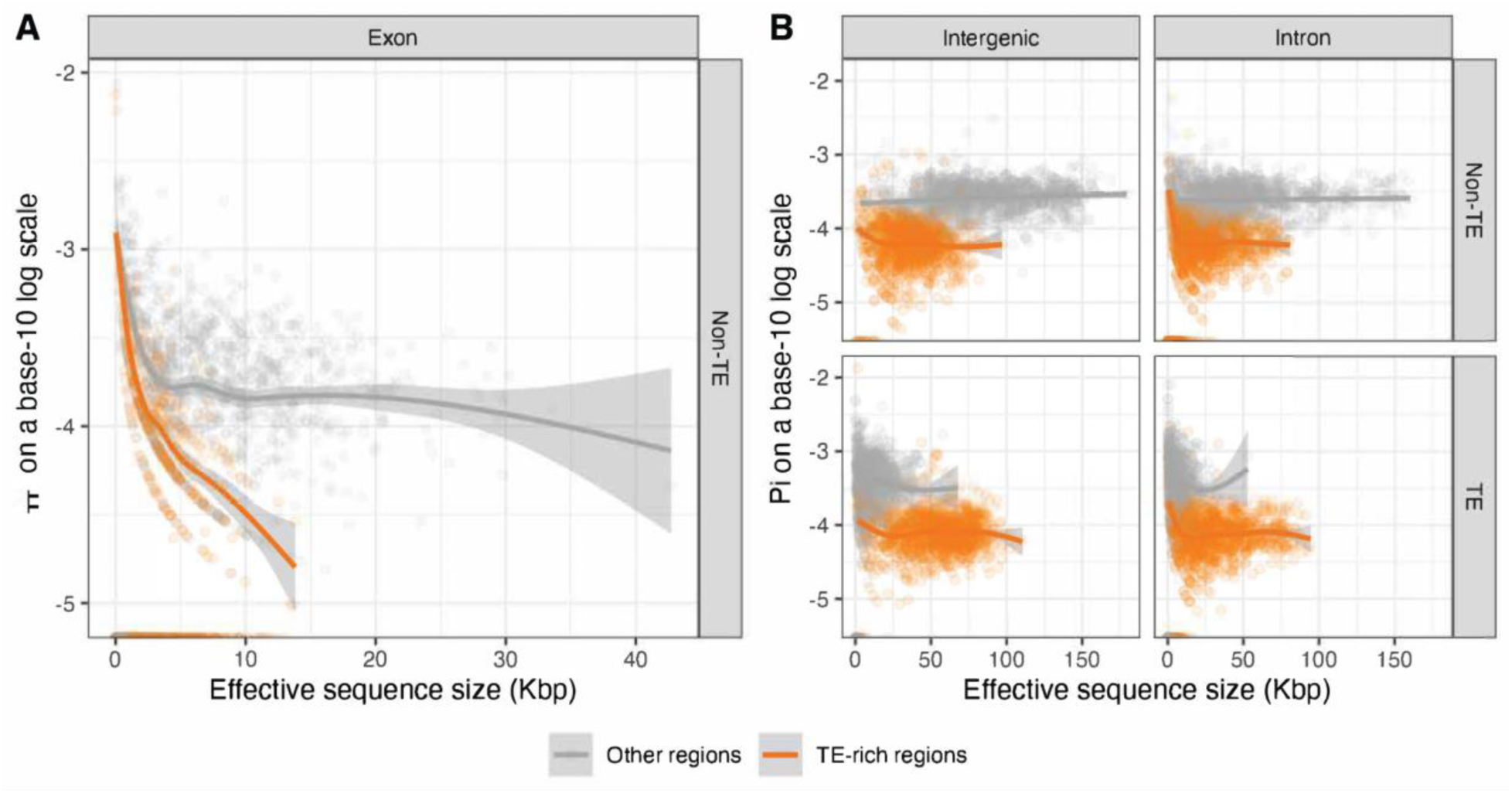
Reduced nucleotide diversity in TE-rich regions. Genomic sequences were first separated into 200-Kbp windows and then separated by gene features (exons, introns and intergenic) and TE annotations (TEs and non-TEs). π was estimated in each window and plotted against their effective sequence sizes, colored by whether the windows were in TE-rich regions. (A) Exons; (B) Intergenic and intronic TEs and non-TEs. Smoothed lines represent conditional means, and the shaded area around the lines correspond to the 95% confidence intervals.

To explore if there were significant differences in large scale patterns of molecular evolution between the two genomic compartments, we identified orthologous genes in *E. verticillata* and calculated synonymous divergence (dS), non-synonymous divergence (dN) and the dN/dS ratio. We found that genes located in TE-rich regions have lower dS (p < 0.0001, Wilcoxon test), but higher dN (p < 0.0001, Wilcoxon test) and dN/dS ratios (p < 0.0001, Wilcoxon test) when compared to genes outside these regions (Supplementary figure 20). This pattern is suggestive of significant differences in selection efficiency between the compartments. To further explore that idea we calculated π at 0-fold (π_0_) and 4-fold (π_4_) degenerate sites. We found that genome-wide π_0_ was higher relative to π_4_ in TE-rich regions (π_0_ = 3.56 × 10⁻⁵; π_4_ = 5.53 × 10⁻⁵) compared to other regions (π_0_ = 7.76 × 10⁻⁵; π_4_ = 3.34 × 10⁻⁴) (Supplementary figure 12A). Further, the contrast of π_0_ and π_4_ is not significant in TE-rich regions (p=0.2744) but was significant in the other region (p<0.0001) (Supplementary figure 12B).

### Lower GC3 is the Major Signature of Genes in TE-rich Regions, not DNA Methylation

To maintain genome integrity, TEs in plants are often silenced through DNA methylation and histone modification (Liu and Zhao 2023). However, the reduced abundance of DNA methylation in insects suggests that DNA methylation might have lost their role in TE silencing (Bewick et al. 2017; Provataris et al. 2018). To explore the potential association between TEs and DNA methylation in fig wasps, we annotated methylated CpG dinucleotides in the genome assembly using the HiFi reads. In total 81K CpG 5mC methylation sites were detected, comprising only 0.6% of the total CpGs in the genome. We found methylation to be significantly lower in the TE-rich region (p<0.001, Wilcoxon test) (Supplementary figure 14). We next examined the distance between each methylated CpG to their nearest genes or TEs. We generated two null distributions by permutating methylated CpGs among all CpGs (labeled “permuted”) and randomly assigning new positions to each methylated CpGs across the whole genome (labeled “random”). Genes were enriched in CpG methylation (Figure 3A) when compared with null distributions, especially from 500 bp upstream and 5000 bp downstream of the start codon (Supplementary figure 15). On the other hand, the average distance between TEs and methylated CpGs is greater than the average distance observed in the null distributions (Figure 3A), likely resulting from the fact that methylation is restricted to genes. Further analyses show that evolutionarily conserved genes are more heavily methylated (Supplementary table 7, Supplementary figure 17), and methylated CpGs were also more enriched in exons than in introns (Supplementary table 8).

**Figure 3.**
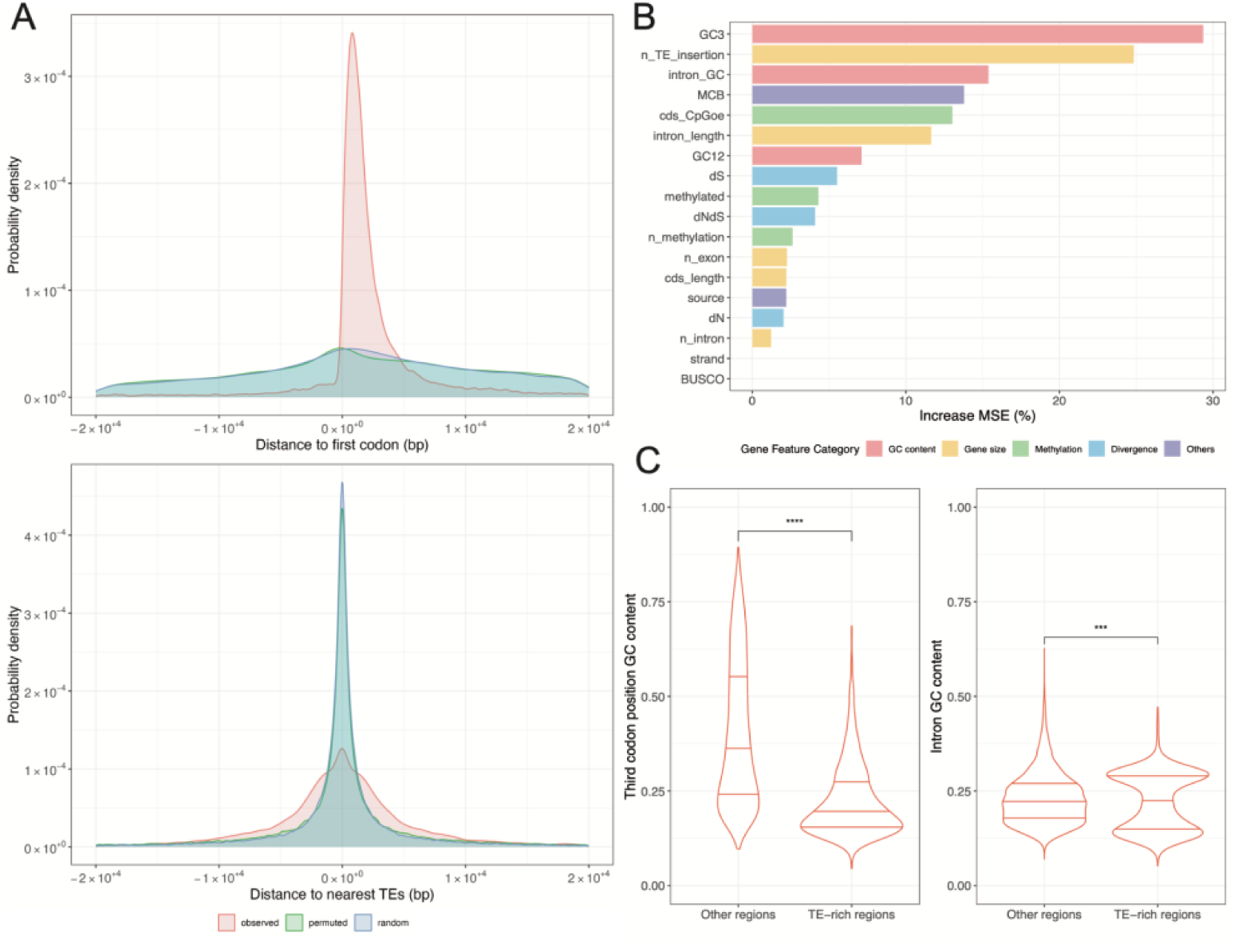
Comparison between genes in TE-rich regions and other regions: (A) Probability density of distances between methylated CpGs to genes and TEs with upstream regions represented by negative coordinates. (B) Importance of gene features in predicting genes in TE-rich regions. Gene features were grouped by biological significance. (C) Comparison of GC3 and intron GC content.

To identify the most critical genomic features associated with genes in TE-rich vs other regions we selected a high confidence set of 6,615 protein coding genes by filtering out genes without orthologs in *E. verticillata* or without introns. A dataset of 18 gene features, organized into five categories was compiled to explore differences between genes located in TE-rich regions and those in other regions. A random forest model was trained to predict whether a gene is in a TE-rich region based on these gene features. The training and testing accuracy reached 82.06% and 78.58%, respectively. Importance analysis revealed that features related to GC content were the main contributors to model accuracy, with GC content at the third codon position (GC3) being the most influential factor (Figure 3B). GC3 content in TE-rich regions was significantly lower than that of genes located on the arms of chromosomes (p < 0.01, Wilcoxon test) (Figure 3C). Additionally, GC3 content in these TE-rich regions was even more reduced compared to the genome-wide GC context. Intron GC content in TE-rich regions showed a bimodal distribution. Specifically, genes with a higher number of TE insertions have a higher intron GC content close to the genome-wide average GC content (∼30%) (Supplementary figure 19), while introns in genes with fewer TE insertions are extremely AT-biased. Furthermore, features associated with gene size, specifically the number of TE insertions and intron length, are also significant features that distinguish genes in TE-rich and other regions. Maximum likelihood codon bias (MCB) was identified as a relevant feature, albeit less impactful. As MCB and GC3 content are highly correlated (Supplementary figure 21), the pattern of MCB is likely driven by GC3 content.

## Discussion

Fig wasps have become a remarkable system to study plant-pollinator mutualisms due to their obligate association with figs (Machado et al. 2001; Herre et al. 2008; Cruaud et al. 2012). However, we have only recently started uncovering the genomic underpinnings of this mutualism, albeit with still limited genomic resources available for most fig wasp genera (Xiao et al. 2013; Cooper et al. 2020; Zhang et al. 2020; Wang et al. 2021; Xiao et al. 2021; Chen et al. 2022). We present the first study exploring the genome organization and evolution of a fig wasp from the Americas, *Pegoscapus hoffmeyeri* Sp. A, the pollinator of *Ficus obtusifolia* in Panama. The genome is 1.32 to 1.86 times larger than the genomes of other fig wasp species sequenced so far and the genome size expansion is the result of increased TE content. The abundance of TEs uncovered here challenges the notion that ecological isolation inhibits TE accumulation by limiting horizontal gene transfer (Gilbert et al. 2021).

### The Genome of *P. hoffmeyeri* sp.A Exhibits Two Distinct Compartments with Large-scale Bimodal TE Enrichment Patterns

TEs exhibit specific distribution patterns throughout eukaryotic genomes (Sigman and Slotkin 2016; Bourque et al. 2018). Intrinsic insertion preferences shape TE-specific localization within or close to some genomic features during the initial integration phase, while selective pressures bias overall TE retention in the post-insertion phase (Sultana et al. 2017). For example, the LTR retrotransposon *Ty1* in yeast preferentially inserts itself upstream of genes transcribed by RNA Polymerase III (Guo et al. 2015; Cheung et al. 2016; Cheung et al. 2018). On the other hand, TEs close to genes or TEs that disrupt exons are more rapidly purged (Cridland et al. 2013; Cridland et al. 2013; Stritt et al. 2018). Given that sequence characteristics only explain the enrichment of few TE families (Liao et al. 2000; Liu et al. 2005; Linheiro and Bergman 2012), Mbp-scale TE enrichment patterns that apply to most TEs in general more likely result from location-based differential selection pressure. In gene-rich euchromatic regions, a stronger selective pressure exists to remove TEs that disrupt exons and regulatory regions, or that introduce repressive epigenetic marks into neighboring genes (Sigman and Slotkin 2016; Huang et al. 2022) and consequently TEs are more likely to be retained in intergenic regions, where other inactive TEs can also serve as a buffer between them and neighboring genes. Given that TEs are also recognized as sources of structural polymorphisms (Gray 2000; Maumus et al. 2015; Mun et al. 2021; Balachandran et al. 2022; Munasinghe et al. 2023; Carpinteyro-Ponce and Machado 2024), purifying selection against TE insertions can also be strong due to their potential to generate deleterious chromosome rearrangements by ectopic recombination between non-allelic TE insertions, posing major risks to genome stability (Kent et al. 2017). However, reduced recombination rates in certain genomic compartments can shelter TEs from purifying selection, promoting their accumulation in these regions (Dolgin and Charlesworth 2008; Sigman and Slotkin 2016). Thus, the interplay between insertion preferences, selection pressures and recombination dynamics can influence the distribution and evolution of TEs within a genome.

In the *P. hoffmeyeri* sp.A genome, we observed a large-scale binary enrichment pattern of TEs, where nearly half of each chromosome, specifically the pericentric region, was populated with TEs. This pattern is consistent with the pericentric heterochromatin observed in *Drosophila* in terms of location and genomic element composition (Hoskins et al. 2015). However, although these regions in the *P. hoffmeyeri* sp.A genome are full of TEs, as TEs are also located in introns, they are still gene-rich as they encompass about 40% of all the annotated protein coding genes in this genome. The TE-rich regions reported here are thus substantially different from known gene-poor and heterochromatic pericentric regions. Since recombination is suppressed in centromeres and pericentromeres (Mancera et al. 2008; Comeron et al. 2012; Nambiar and Smith 2016; Kent et al. 2017), the significant enrichment of TEs in these large pericentric regions is likely the result of reduced recombination (see below).

### GC3 Patterns Provide Indirect but Strong Evidence of Lower Recombination Rates in TE-rich Regions

GC3 content was the most significant feature (of 18 gene features) distinguishing TE-rich regions from other regions (Figure 3B). GC3 is significantly lower in TE-rich regions and this effect is more pronounced than metrics that are intuitively correlated with TEs, such as the number of TE insertions in introns (Supplementary figure 20). It is well-established that lower GC3 content is associated with lower recombination rates through GC-biased gene conversion (gBGC) in mammals (Eyre-Walker 1997; Galtier et al. 2001; Duret and Galtier 2009), other eukaryotes (Pessia et al. 2012), and, more relevantly, in honey bees (Kent et al. 2012). This is because in addition to crossover, recombination can also lead to gene conversion, which could favor G/C alleles over A/T alleles, resulting in an increase of GC content following conversion. Therefore, the observed lower GC3 content in TE-rich regions could be a result of the absence of gBGC that recovers the GC content induced by AT biased mutations, as recombination rates are lower in these regions.

### Patterns of Nucleotide Diversity and Gene Evolution in TE-rich regions

Nucleotide diversity is 75% lower in pericentric TE-rich regions (Figures 1C, 2). This pattern was consistent across exonic, intronic and intergenic TE and non-TE sequences, and is also robust against artifacts such as mapping rates and effective sequence sizes. In essence, we found that TE enrichment is associated with reduced nucleotide diversity in neighboring sequences. This large-scale pattern can only be explained by reduction in recombination rates and/or mutation rates as opposed to demographic histories, which are expected to affect the whole genome. Although lower mutation rates are consistent with the lower dS observed in genes from TE-rich regions, this parameter is not a consistent predictor of these genes in our machine learning model (Figure 3B) and thus provides very weak support for any mutation rate variation among the two genome compartments. Instead, the observed differences in GC3 content support the conclusion that low nucleotide diversity in TE-rich regions is likely the result of enhanced Hill–Robertson interference (HRI) (Hill and Robertson 1966) caused by reduced recombination rates. Given that TE-rich regions harbor a substantial number of genes, and that most of those genes should be under some level of purifying selection, HRI effects are likely to be strong in these regions due to reduced recombination.

Genes located in TE-rich regions had higher dN/dS ratios when compared to genes outside these regions (Supplementary figure 20). At first glance, this is consistent with patterns previously observed in TE islands from inbreeding ants where genes in TE-rich regions exhibit higher adaptive divergence rates (Schrader et al. 2014). In this genome, however, we do not interpret higher dN/dS as an indicator of faster adaptation. As shown in avian and mammalian species, the absence of recombination and gBGC is associated with an increase in dN/dS ratios (Rousselle et al. 2018) due to the reduced efficiency of purifying selection when HRI effects are strong (Rousselle et al. 2018), leading to an inflation of dN values. Instead of adaptive evolution, genes in these low recombination regions are more likely accumulating deleterious mutations, an effect known as Muller’s ratchet (Muller 1964; Felsenstein 1974). This is consistent with our observation that the ratio of π at zero-fold degenerate sites to four-fold degenerate sites is higher in TE-rich regions compared to other regions (Supplementary figure 12). Furthermore, a decrease in gBGC that results in lower GC content may also slightly decrease mutation rates because cytosine and guanine exhibit higher inherent mutability (Supplementary figure 22), which consequently contributes to lower rates of synonymous divergence. Therefore, the higher dN/dS is likely a combination of direct and indirect consequences of low recombination rates in the TE-rich regions.

### Genes in TE-rich Regions may Evolve at the Epigenetic Level

If DNA methylation suppresses TE activity, as observed in plants (Sigman and Slotkin 2016), we should see enrichment of methylated CpGs in TEs. Instead, our analysis on the distance between methylated CpGs to genes and TEs found enrichment of methylated CpGs only in gene bodies (Figure 3A) and there is no direct association between DNA methylation and TEs. We also found methylated genes are more likely to be evolutionarily conserved BUSCO genes (Supplementary figure 17, Supplementary table 7), consistent with the idea that the function of DNA methylation in insects is to regulate the expression of genes involved in basic biological processes (Elango et al. 2009; Foret et al. 2009; Sarda et al. 2012; Glastad et al. 2014). Given that genes located in TE-rich regions are also more heavily methylated (Supplementary figure 20), it is likely that they could be evolving at the epigenetic level (Schmitz et al. 2011; Hagmann et al. 2015; Torres-Garcia et al. 2020; Ashe et al. 2021; Sarkies 2023; Wilson et al. 2023). In the *P. hoffmeyeri* sp. A genome, nearly half of the annotated genes, including 30% of the BUSCO genes, are located in these TE-rich regions, suggesting that gene expression does occur in TE-rich regions. As many of these genes have gone through dramatic intron sequence elongation due to TE insertions (Supplementary figure 6), selection may favor increased methylation as a compensatory mechanism necessary for proper gene expression (Jeong et al. 2018; Wu et al. 2022). Given that DNA methylation and histone modifications can be highly related (Hunt et al. 2013), these genes could also adopt histone modifications to facilitate expression within the TE-rich context (Saha and Mishra 2019). For example, although H3K9me3 is a conserved repressive epigenetic modification that induces chromatin compaction by recruiting heterochromatin protein 1 (HP1), it is also required for the expression of heterochromatic genes and suppression of aberrant genes (Ninova et al. 2020). Similar molecular mechanisms are likely being adopted during evolution to ensure gene expression.

### Implications of Genome Size Expansion on Fig Wasp Evolution

The genome sizes of most hymenopteran species sequenced so far range from 180 to 340 Mbp, with approximately 12,000 to 20,000 genes (Branstetter et al. 2018). Furthermore, they have fairly low GC content (30-45%) (Branstetter et al. 2018) and low TE content relative to other arthropods (Petersen et al. 2019). Although *P. hoffmeyeri* sp. A has a very large genome compared to other fig wasps, the TE-free size of its genome is similar to that of other fig wasps (Table 1). This suggests that TEs play a major role in the evolution of genome size in this group of insects. In contrast to the consistent and moderate TE activity inferred from the TE landscape of *E. verticillata*, *P. hoffmeyeri* sp. A has experienced an intense burst of TE activity that peaked approximately 5.6 mya. As a consequence of the absence of alignable TE copies when they are too degraded to be recognized, the TE landscape can only show TE activities up 10 to 15 mya (40-60 Kimura divergence), which is more recent than the crown ages of every pollinating fig wasp genera (Cruaud et al. 2012). Thus, the TE landscapes shown for *P. hoffmeyeri* sp. A and *E. verticillata* are independent and represent a partial TE activity history of each genus and some species-specific TE activities. To gain a comprehensive understanding of when the TE burst started in *Pegoscapus* and what TEs contributed to the genome size expansion, a more extensive sampling of high-quality fig wasp genomes across multiple genera will be necessary.

There could be ecological reasons behind the observed genome size expansion. The adaptive hypothesis of genome size evolution posits that a larger genome size may provide evolutionary advantages, since organisms may benefit from larger cells and body sizes (Gregory et al. 2000). However, comparative studies have failed to establish a correlation between body size and genome size (Xu et al. 2021; Yuan et al. 2021). In the case of fig wasps, an increase in body size may enhance dispersal capabilities that facilitate access to receptive syconia. However, it can also interfere with their capacity to enter the fig syconium through the narrow ostiole (Liu et al. 2013). Notably, even though *P. hoffmeyeri* sp.A is one of the largest pollinator species in Panama (Herre 1989), its female body size (2mm) (Wiebes 1995) is still similar to that of *E. verticillata* (Waterston 1921). Although fig wasp body size is likely the result of coevolution with the size of their host fig ostiole (Weiblen 2002), it will be interesting to determine if there is a significant correlation between body size and genome size in genus *Pegoscapus*.

## Acknowledgements

This study was funded by National Science Foundation grants MCB-1716532 and DEB-2225083 to C.A.M. We thank Thomas Kocher, Phillip Johnson and Steve Mount for helpful discussions and comments on the manuscript. We are grateful to the Institute for Genome Sciences (University of Maryland) for generating the sequencing data.

## Data Availability

All scripts were deposited at the project github repository (https://github.com/ZexuanZhao/Pegoscapus-hoffmeyeri-sp.A-genome-paper). PacBio HiFi reads and the genome assembly of *P. hoffmeyeri* sp. A are available in NCBI under project number PRJNA1196887. Illumina short reads were deposited under project number PRJNA1198996.

