## Supplementary Materials for "Evolutionary Consequences of Unusually Large Pericentric TE-rich Regions in the Genome of a Neotropical Fig Wasp"

**for**

### Supplementary Methods

#### Mutational Spectrum Analysis

We inferred the mutational spectrum from individual genotypes called from Illumina reads. We first filtered out genotype calls with a genotype quality (GQ) of less than 30, and then removed loci with more than 1 missing individual. Assuming a parsimonious scenario where each segregating allele was the result of a single mutation from the major allele to the minor allele, we then counted the number of mutations with directionality among the biallelic loci whose minor allele frequencies were lower than 30%. We then separated these counts based on their TE annotations (TE or non-TE) and their regions (TE-rich regions and other regions).

#### Zero-Inflated Poisson Regression Analysis of π at 0-fold vs 4-fold Degenerate Sites in TE-rich Regions and Other Regions.

We used a zero-inflated Poisson regression model to investigate how sites (0-fold and 4-fold degenerate sites in coding regions) and regions (TE-rich regions and other regions) affect π. The genome was first partitioned into 1-Mbp windows and π per window was calculated using pixy v1.2.7.beta1 as the total number of pairwise nucleotide differences divided by the total number of comparisons performed separately at 0-fold vs 4-fold degenerate sites. Windows were assigned to TE-rich regions or other regions based on our previous classification. As π is essentially counts of differences normalized by the sequence length, π was scaled up by $10^6$ and rounded to ensure compatibility with the Poisson distribution.

A zero-inflated Poisson model was fitted using the `zeroinfl` function from the pscl package v1.5.9 (Jackman 2024) in R. This model accounts for the presence of windows with minimum nucleotide diversity in coding sequences potentially due to recent selection. Estimated marginal means (EMMs) for each site type within each region were calculated using the `emmeans` function from the emmeans package v.1.11 (Lenth 2025). Finally, contrast of the EMMs between $\pi$ at 0-fold and 4-fold degenerate sites were performed to assess statistical significance within each region.

### Supplementary Figures

#### Supplementary Figure 1

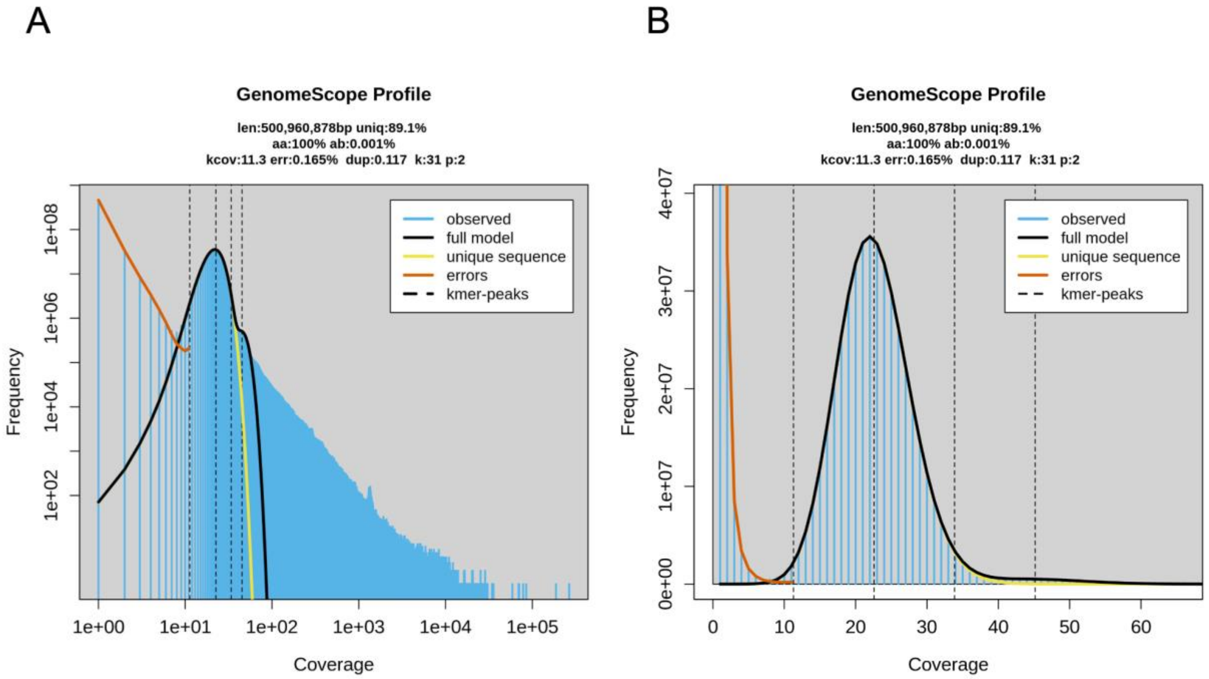


**Supplementary Figure 1: GenomeScope 31-mer Profile of *P. hoffmeyeri* sp. A HiFi Reads:** (A) the coverage histogram on a log scale that demonstrates an over-abundance of high-coverage 31-mers with coverage greater than 100, suggesting a high level of repetitive sequences in the genome. (B) the coverage histogram on a linear scale that is unimodal, indicating the low genetic differences between pooled individuals. Dash lines: expected haploid/diploid/triploid/quadraploid sequencing depth. Len: estimated genome size; uniq: percent of unique sequences; aa: percent of homozygous sites; ab: percent of heterozygous sites; kcov: mean 31-mer coverage of heterozygous sites; err: estimated sequencing error rate; dup: average rate of read duplications; k: kmer size; p: estimated ploidy.

**Supplementary Figure 2**

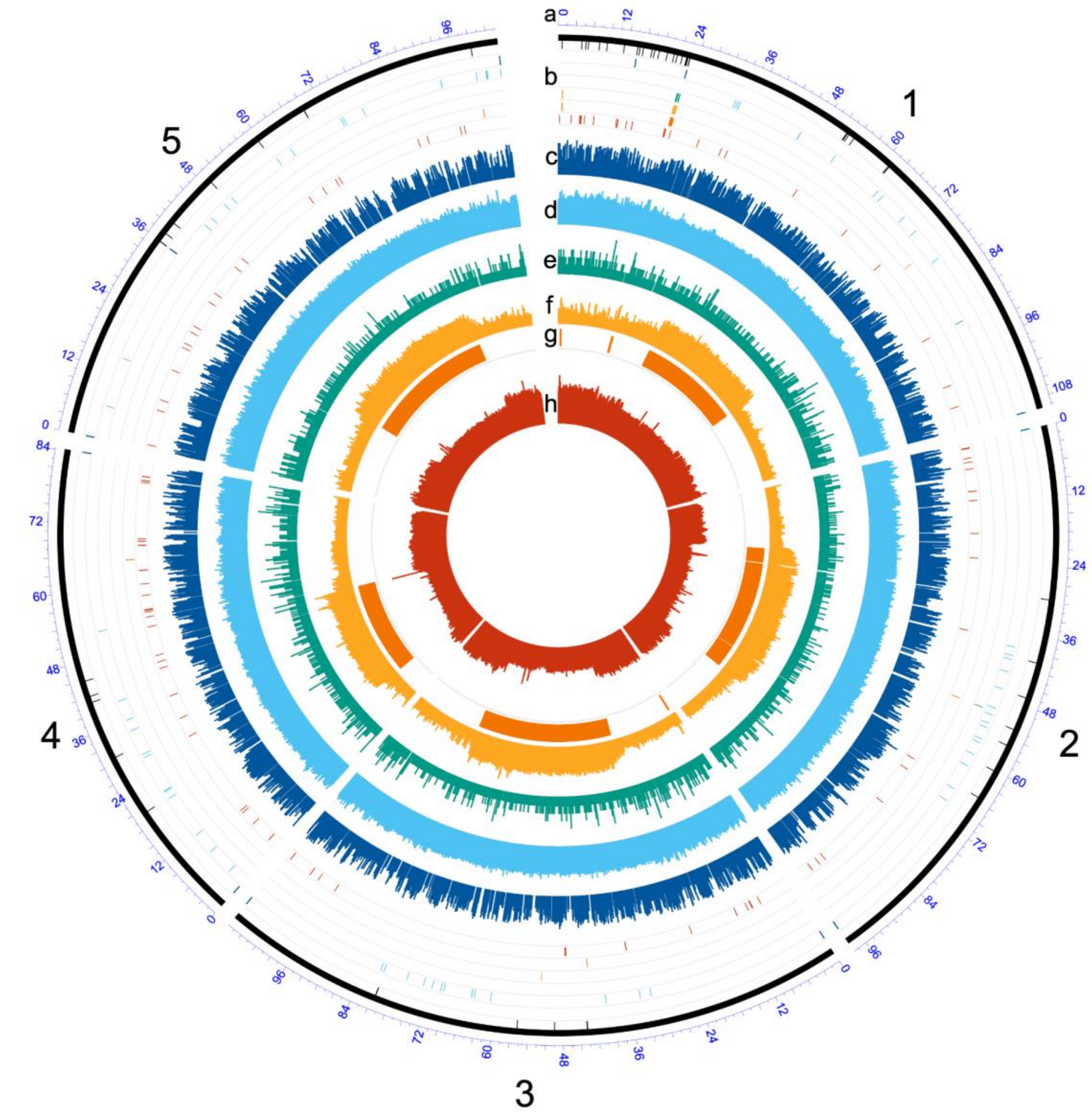


**Supplementary Figure 2 Circos plot of the genome assembly and annotation.** (a) five chromosomal scaffolds; (b) locations of genomic elements from outer to inner circles: contig boundaries in the scaffolds, telomeric repeats, 5S rDNA, 5.8S rDNA, 18S rDNA, 28S rDNA, tDNA; (c) methylation ratio on a base-10 log scale; (d) G+C content; (e) gene density (number of genes per 10-Kbp window); (f) TE density (percent base pairs annotated as TEs); (g) TE-rich region classified by HMM; (h) genome-wide nucleotide diversity on a log-10 scale.

#### Supplementary Figure 3

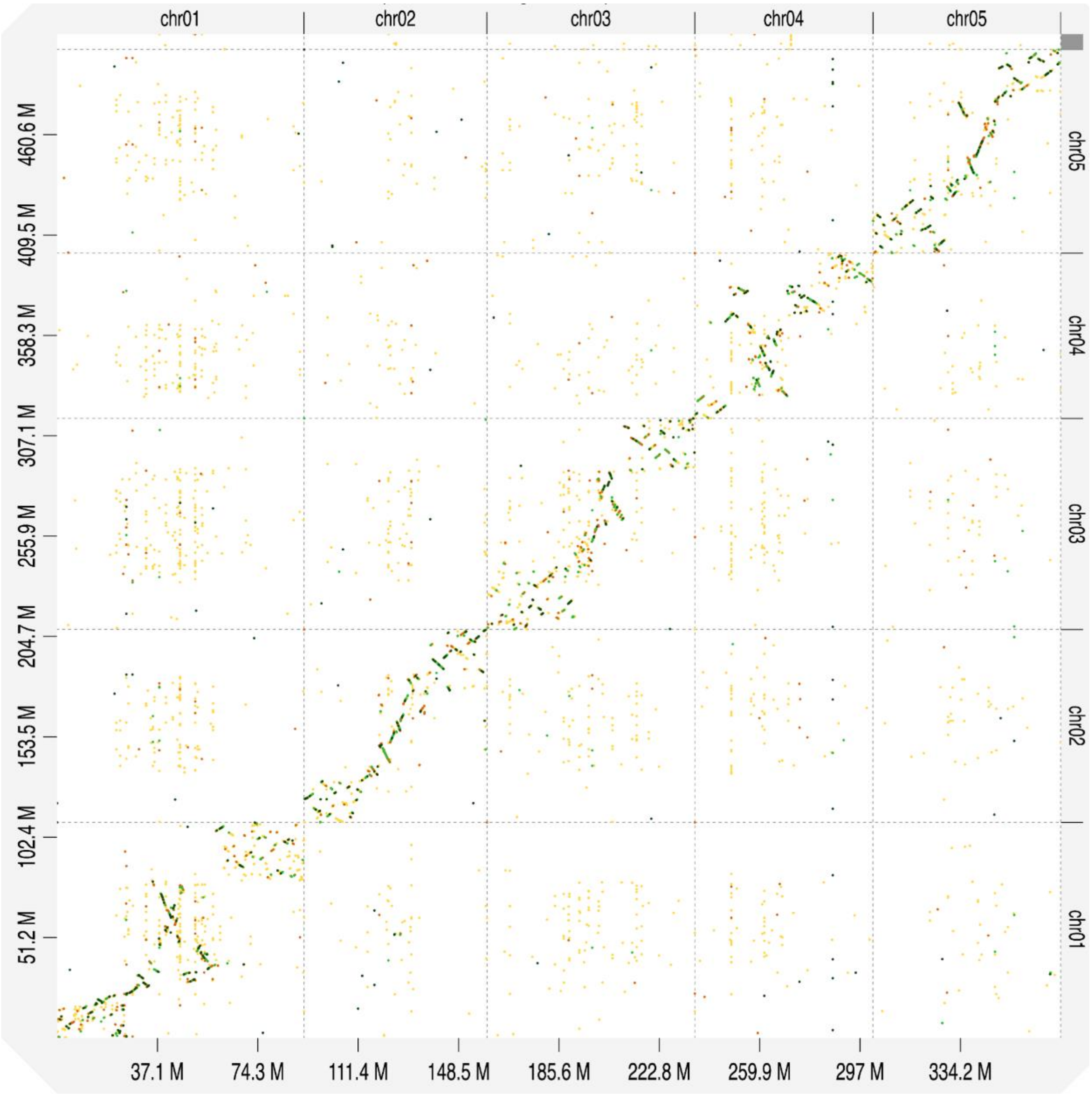


**Supplementary Figure 3 Genome Alignment Dot Plot:** The genome assembly of *P. hoffmeyri* sp. A on the y-axis aligning with the chromosome level assembly of *E. verticillata* on the x-axis.

**Supplementary Figure 4**

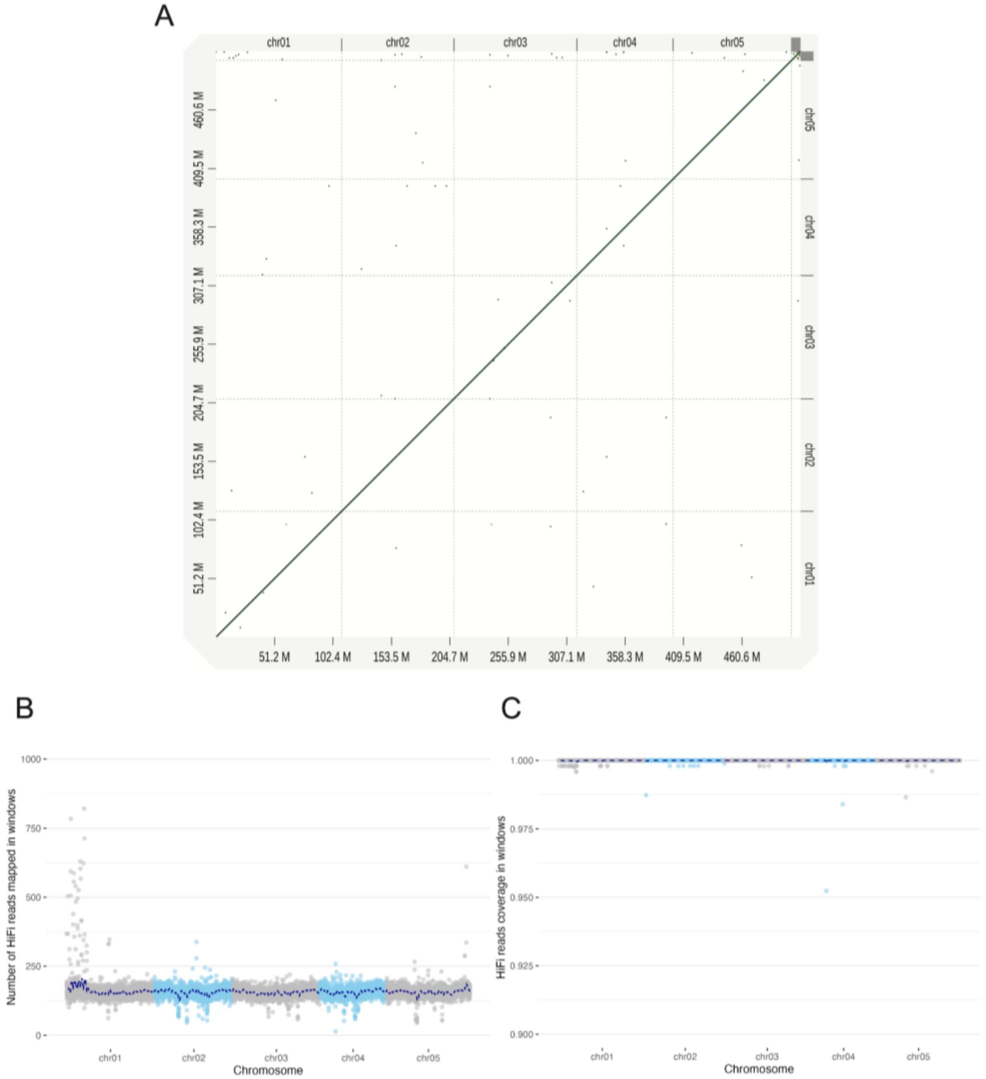


**Supplementary Figure 4 Quality Control for Under-Purging and Over-Purging:** (A) the self-alignment of the *P. hoffmeyri* sp. A genome assembly demonstrates the absence of duplicated sequences as off-diagonal lines. (B) the number of mapped reads in 50-kb non-overlapping windows across the genome, shows no systematically elevated mapping depth, a sign of over-purging.(C) Coverage of mapped read (the percentage of sequences in a window is mapped by at least one reads) shows most positions in the assembly is supported by reads.

#### Supplementary Figure 5

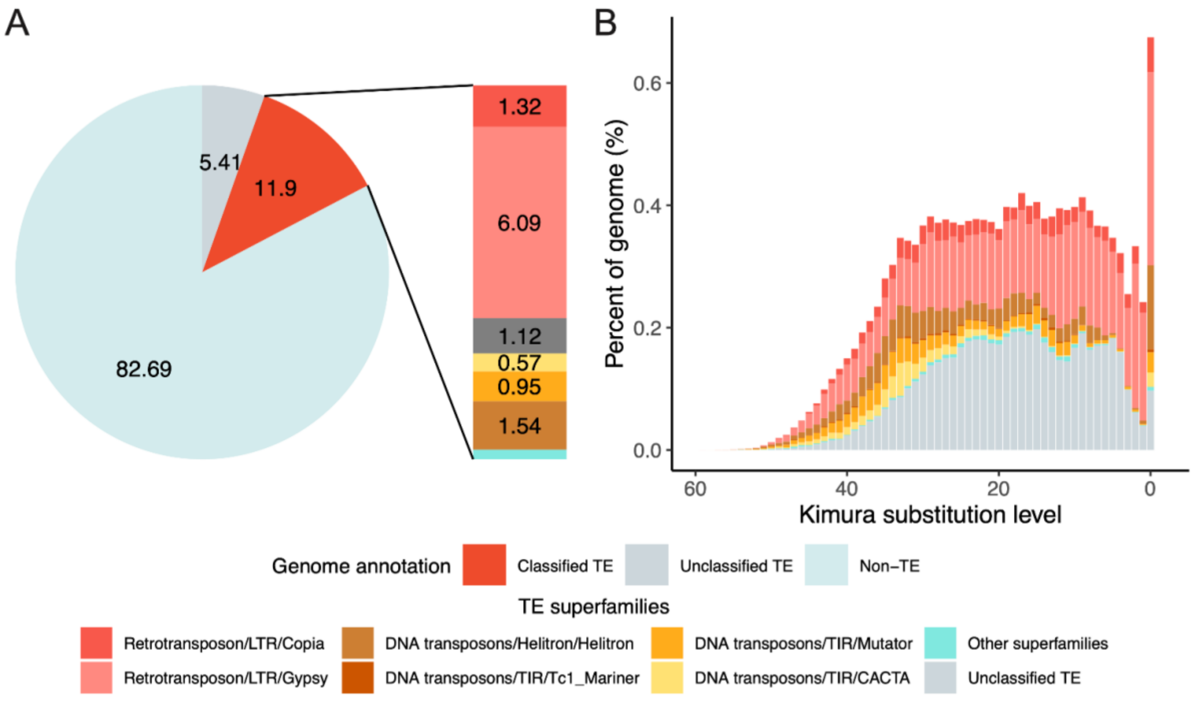


**Supplementary Figure 5 TE Annotation and Landscape of *E. verticillata*:** (A) the percentage composition of the TE annotation, with each component and its corresponding superfamilies labeled. (B) the TE landscape depicting the correlation between the age of TEs and their genome coverage. The X-axis displays the pairwise CpG-corrected Kimura 2-parameter divergence of the TEs relative to the consensus sequence of their respective families, analogous to the age of the TEs. The Y-axis is the percentage of the genome occupied by the TEs.

**Supplementary Figure 6**

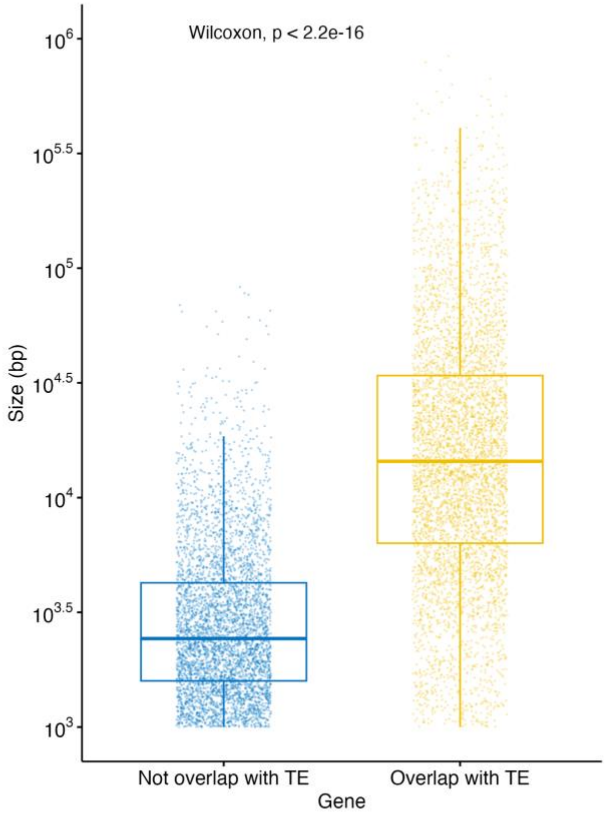


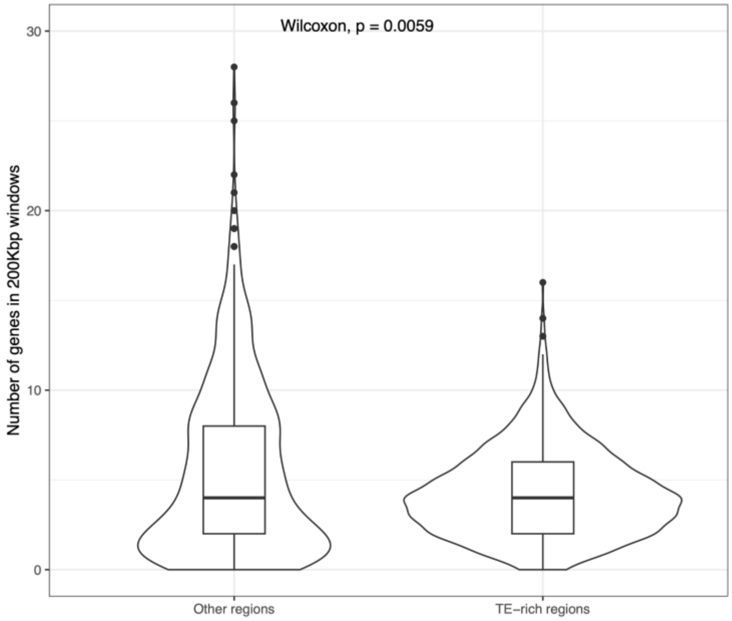


**Supplementary Figure 6: Comparison of Gene Sizes (Top) and Gene Densities (Bottom)**

#### Supplementary Figure 7

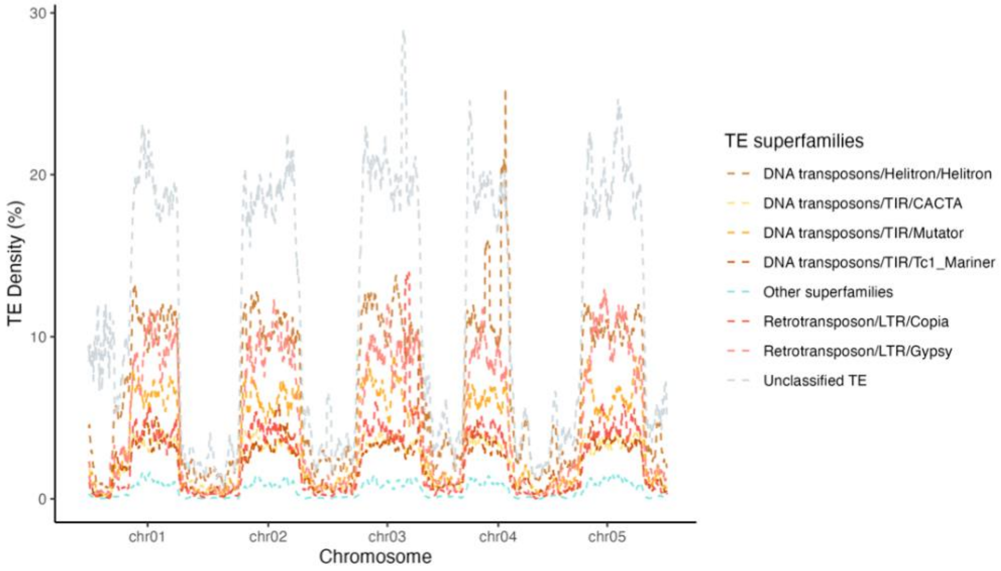


**Supplementary Figure 7 TE Densities per Superfamily Along the Genome.** TE densities separated by superfamilies are shown in non-overlapping 20 Kbp windows along the genome.

#### Supplementary Figure 8

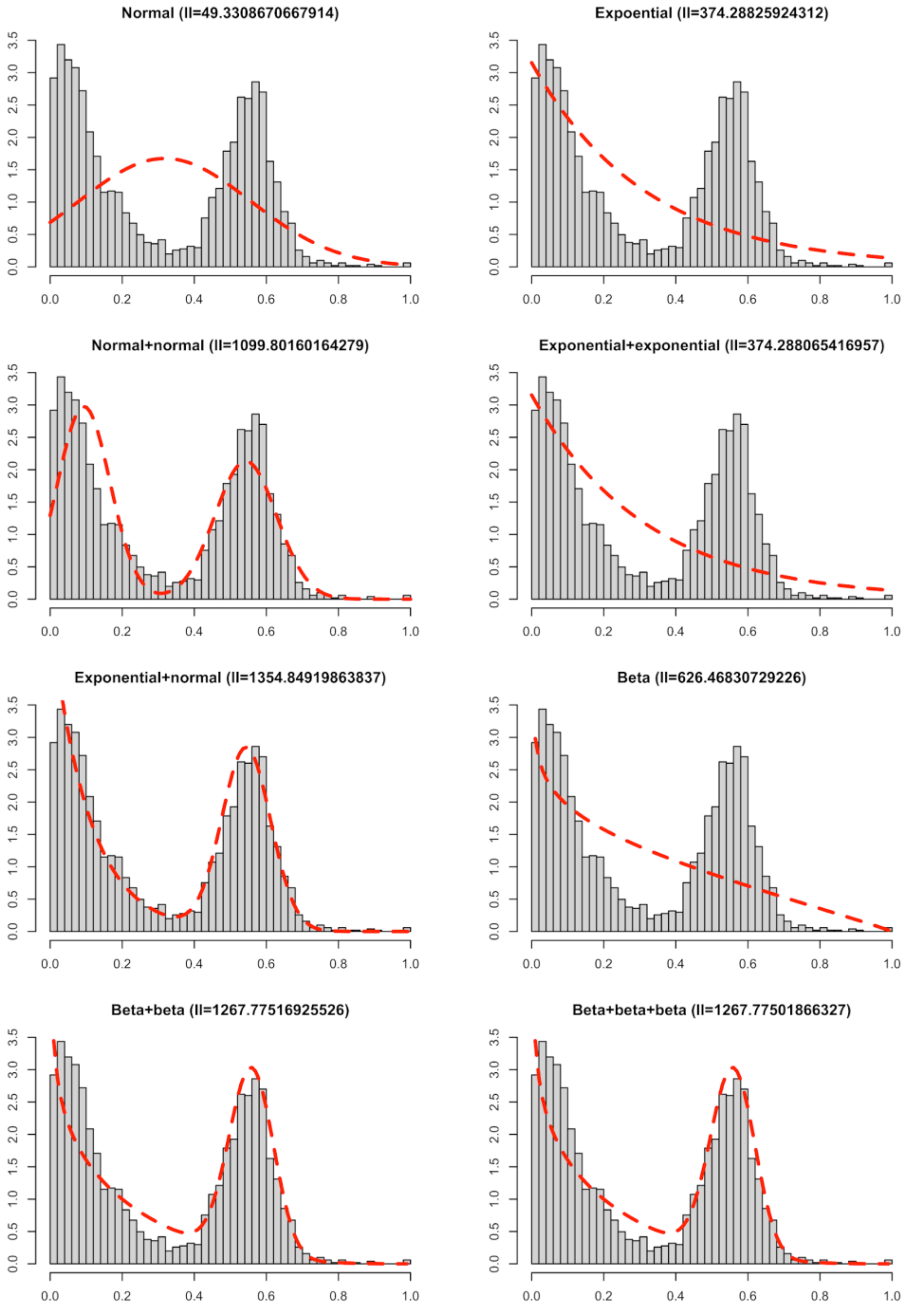


**Supplementary Figure 8 Mixture Distribution of TE Densities.** The histogram of TE densities calculated in 200 Kbp non-overlapping windows were fitted by different mixture distributions. The red dotted lines represent the corresponding likelihood of the maximum-likelihood mixture distributions. See supplementary table 2 for model selection. ll: log likelihood of maximum-likelihood mixture distribution.

**Supplementary Figure 9**

A

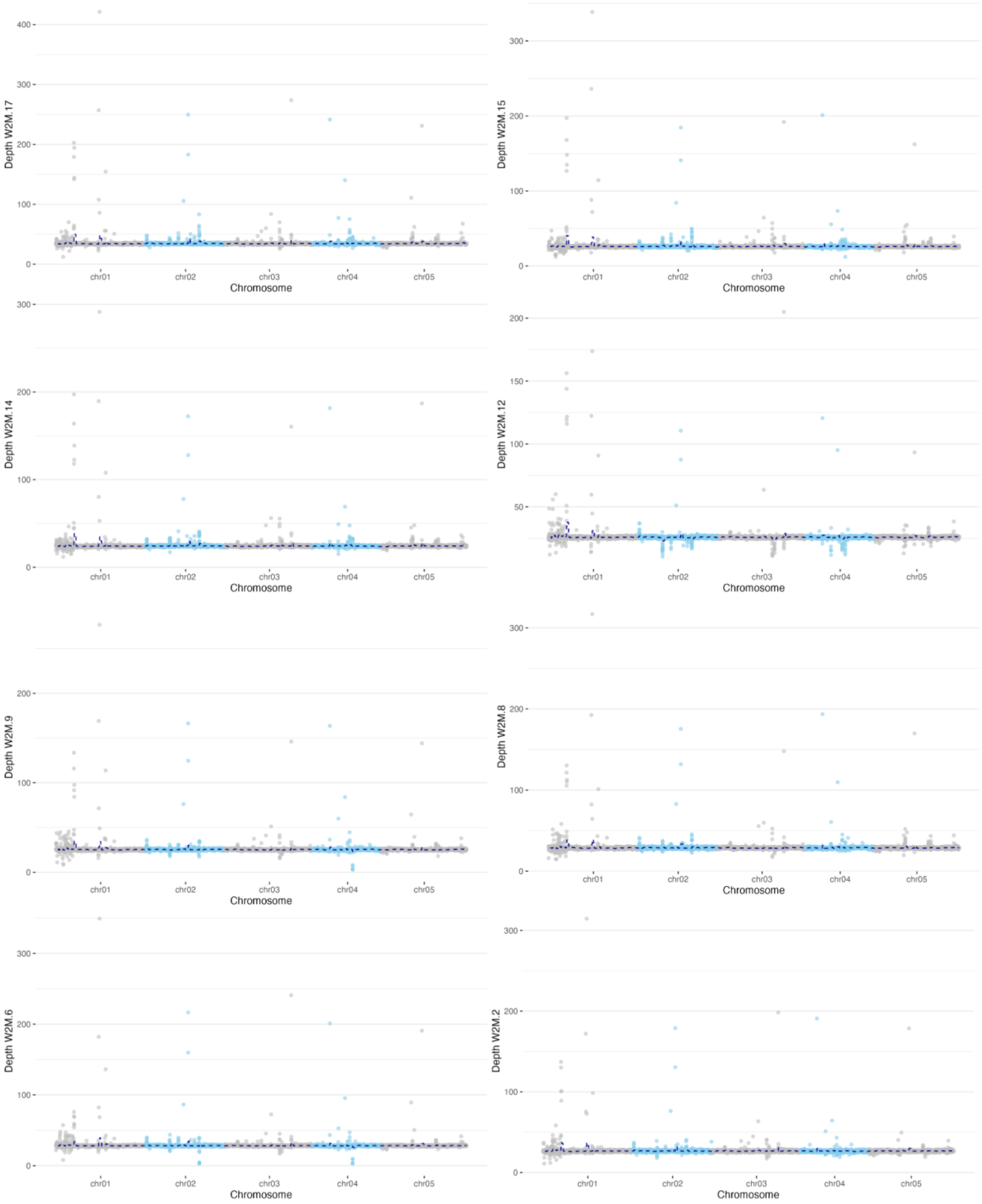

B

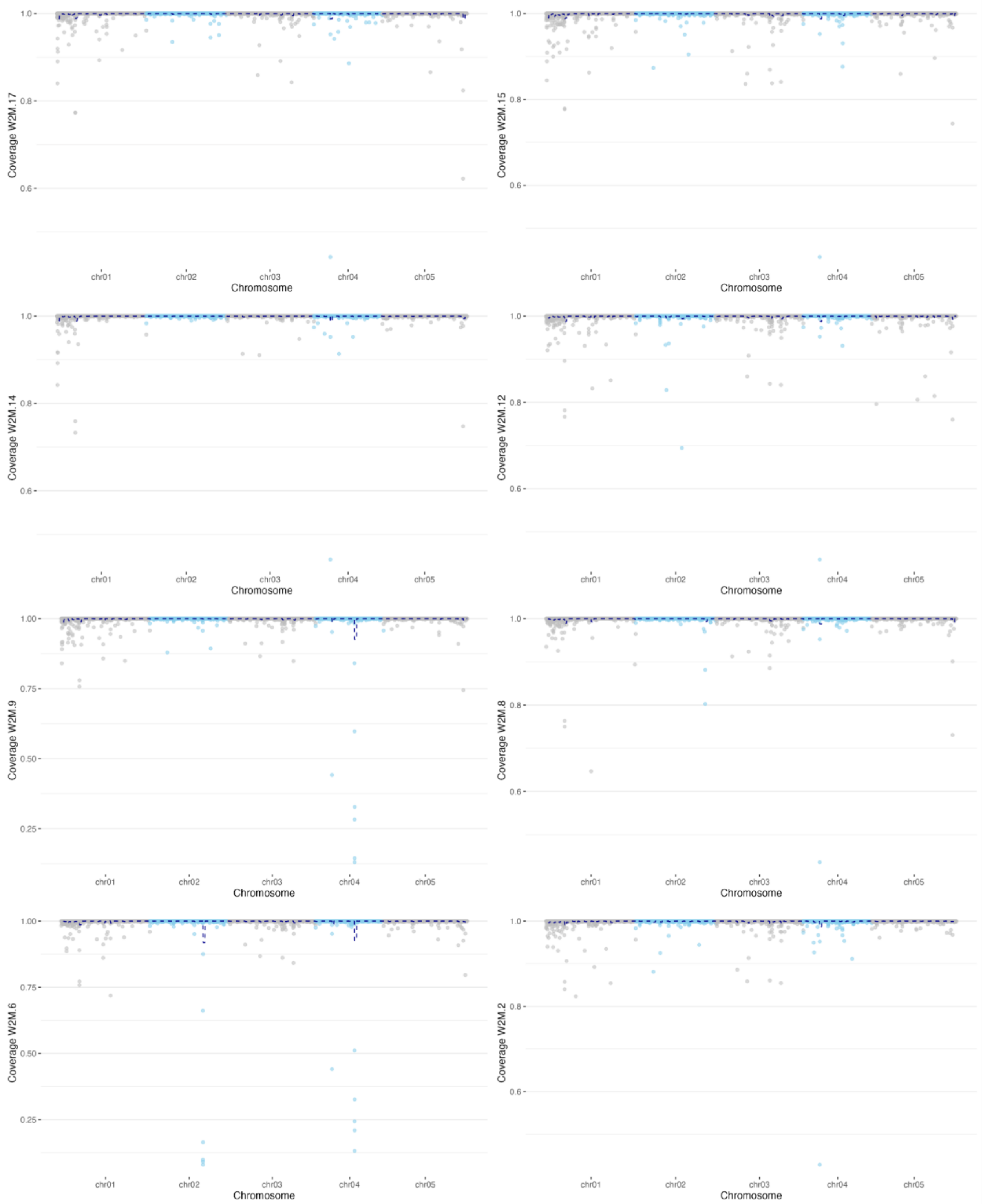


**Supplementary Figure 9 Quality Assessment of Illumina Sequencing of 8 Female Fig Wasps.** (A) mapping depth (B) mapping coverage. Mapping depth: the number of reads mapped to a region. Mapping coverage: the proportion of base pairs in the reference region that is mapped by at least one read.

#### Supplementary Figure 10

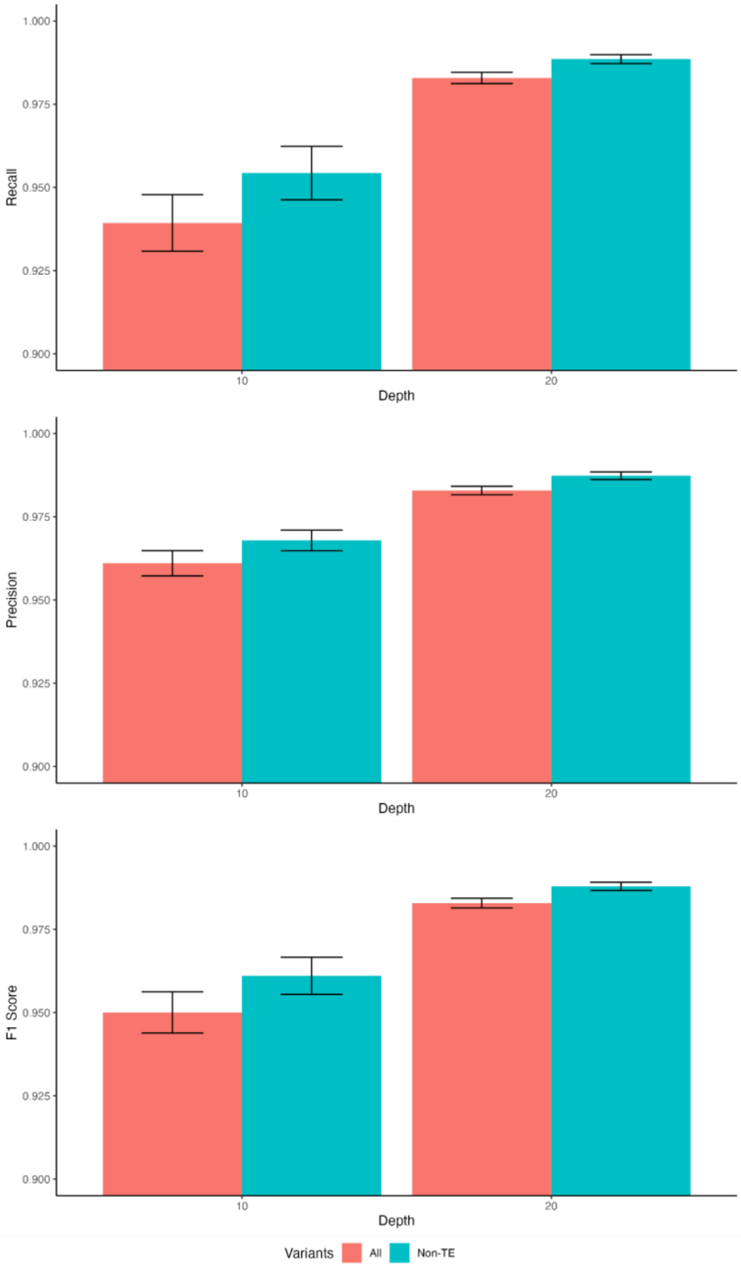


**Supplementary Figure 10 Quality Assessment of Genotyping by Down-sampling:** Evaluation of the performance of DeepVariant at 10x and 20x Sequencing depths compared to all reads. These charts present the recall, precision, and F1 score metrics to assess the genotyping pipeline's performance. Recall, or sensitivity, is calculated as TP / (TP + FN), indicating the genotyper's ability to correctly identify variants. Precision, or specificity, is computed as TP / (TP + FP), reflecting the genotyper's accuracy in variant calling. F1_Score, the harmonic mean of precision and recall, provides a balanced summary of the genotyper's overall performance."

**Supplementary Figure 11**

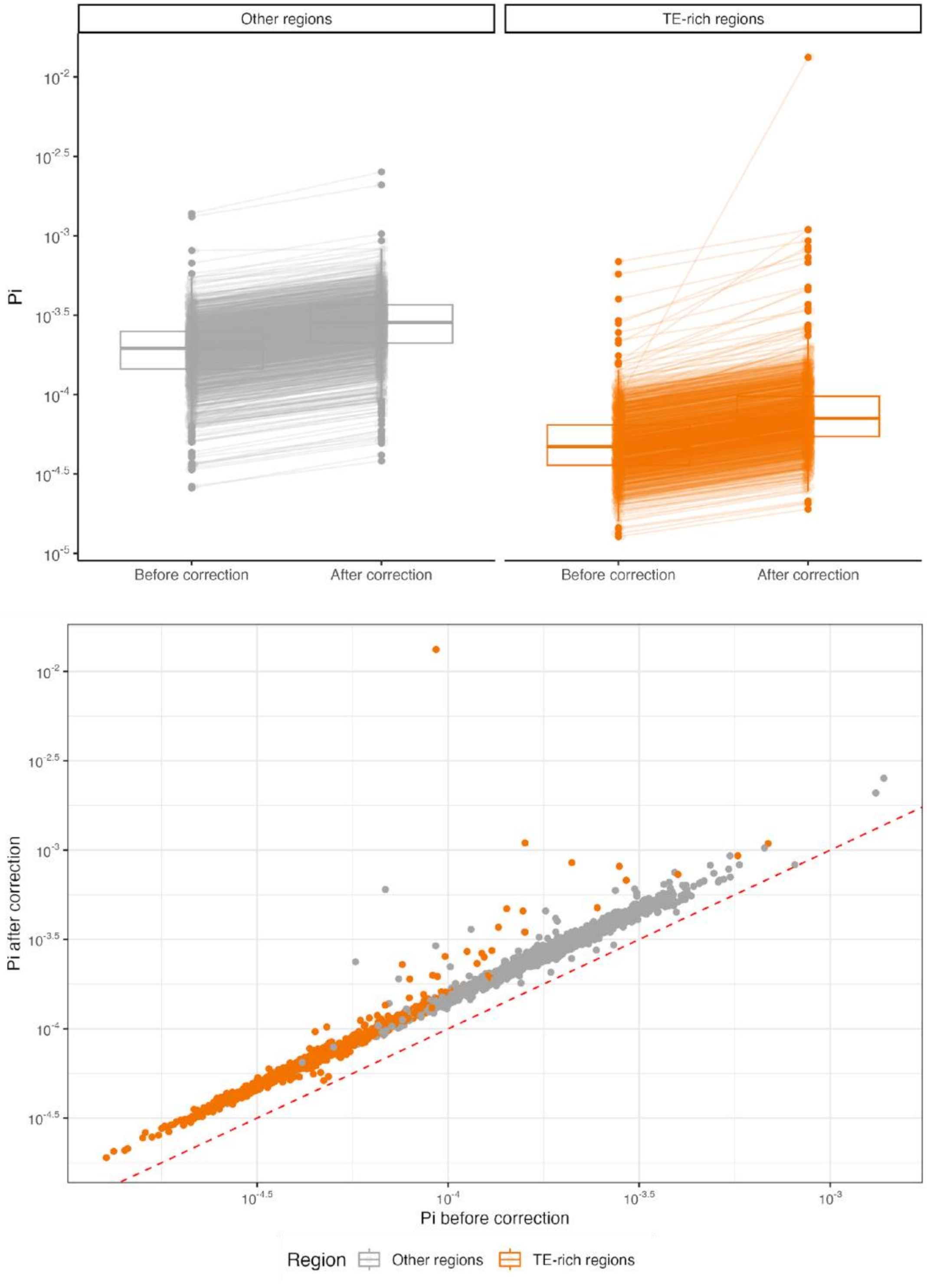


**Supplementary Figure 11 Comparison of Nucleotide Diversity Calculated before and after Missing-Data Correction.** The boxplot shows the significant difference of uncorrected and corrected Pi by considering missing data in TE-dense and background regions. The scatter plot shows the strong correlation on a log-log scale. The red line represents the equation y=x as a reference.

#### Supplementary Figure 12

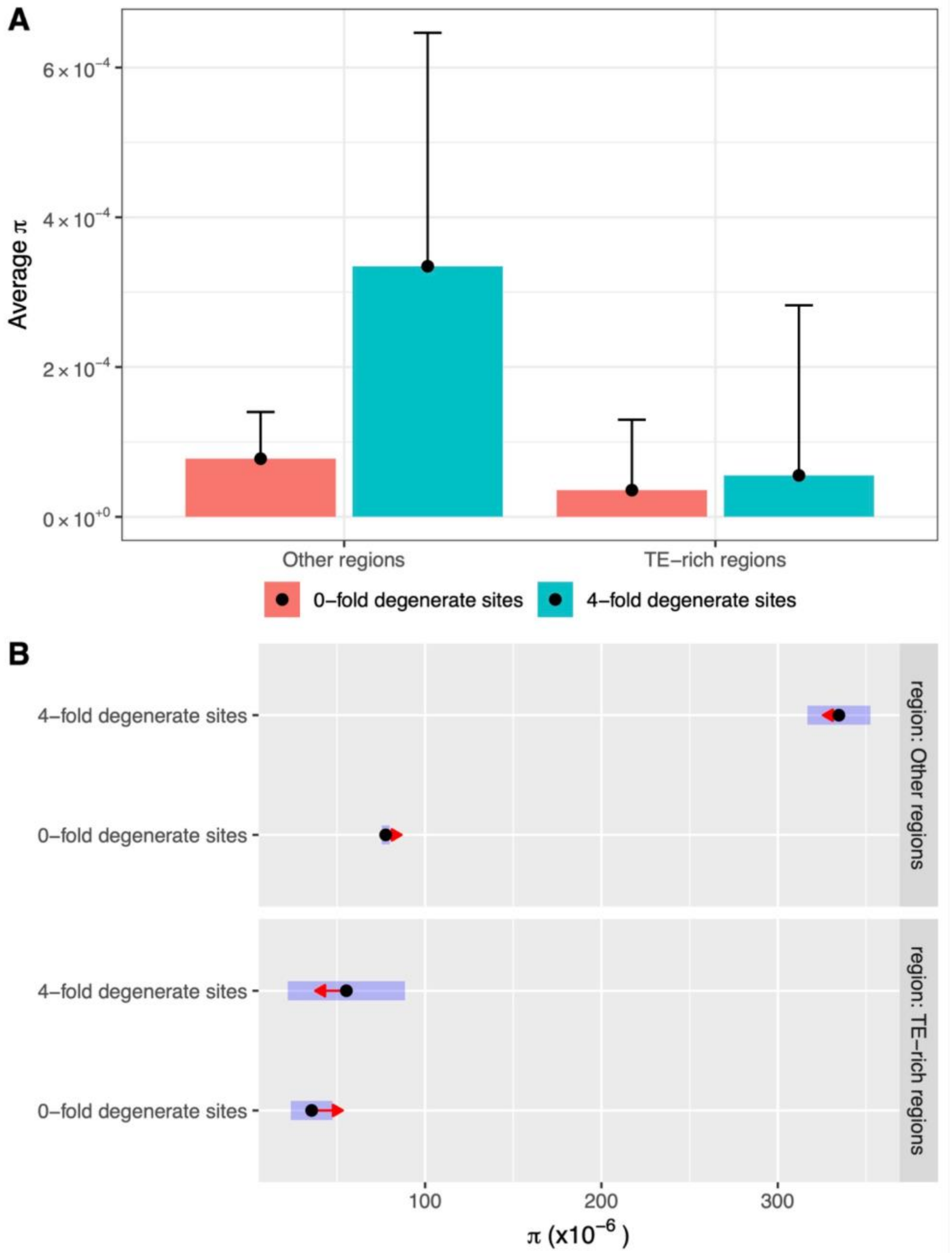


**Supplementary Figure 12 Comparison of Average Nucleotide Diversity at 0-fold vs 4-fold Degenerate Sites in TE-rich Regions and Other Regions.** A) Average π across all loci. Error bars represent the standard deviations calculated from the average π of each category in 1-Mbp windows. B) Marginal means of π estimated by the zero-inflated Poisson regression are plotted as dots. Shaded area represents confidence intervals, and comparison arrows indicate contrast between 0-fold vs 4-fold degenerate sites in TE-rich regions and other regions respectively.

**Supplementary Figure 13**

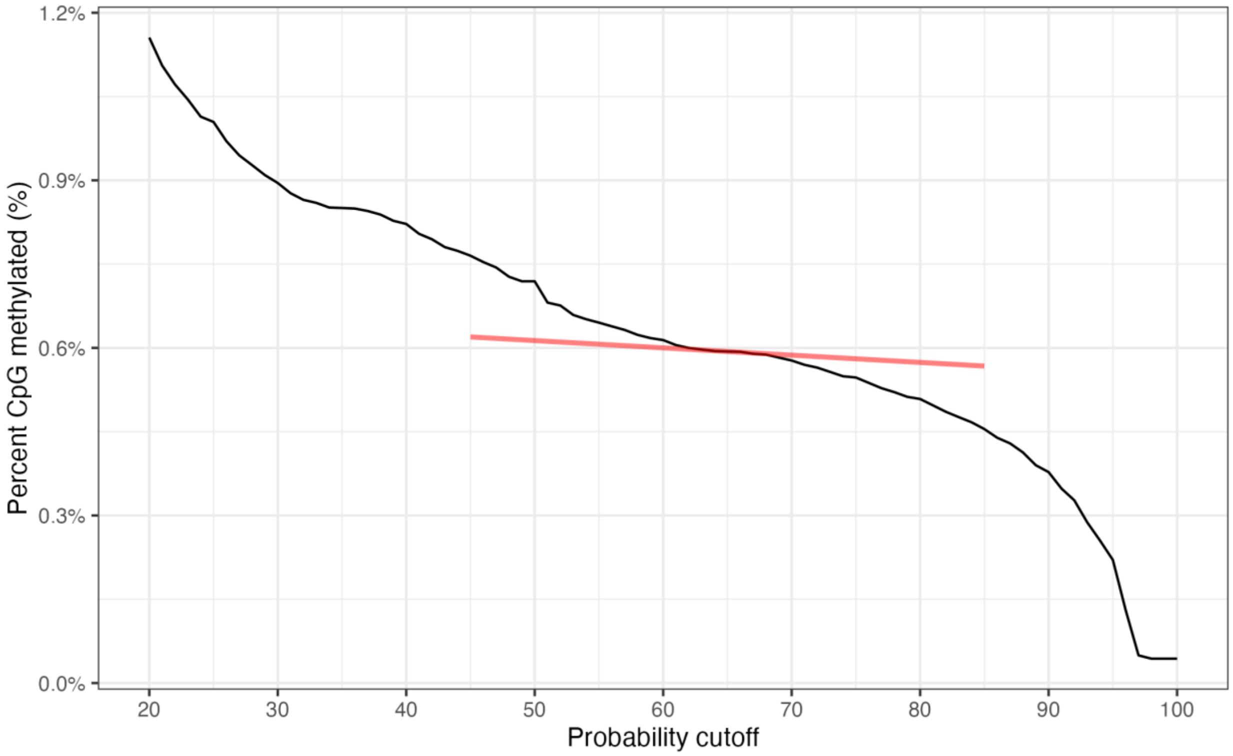


**Supplementary Figure 13: Probability Cutoff versus Percent CpG Methylated.** Red line shows that at probability cutoff = 0.65, the absolute empirical slope is lowest, which should yield the most robust estimate given noise.

#### Supplementary Figure 14

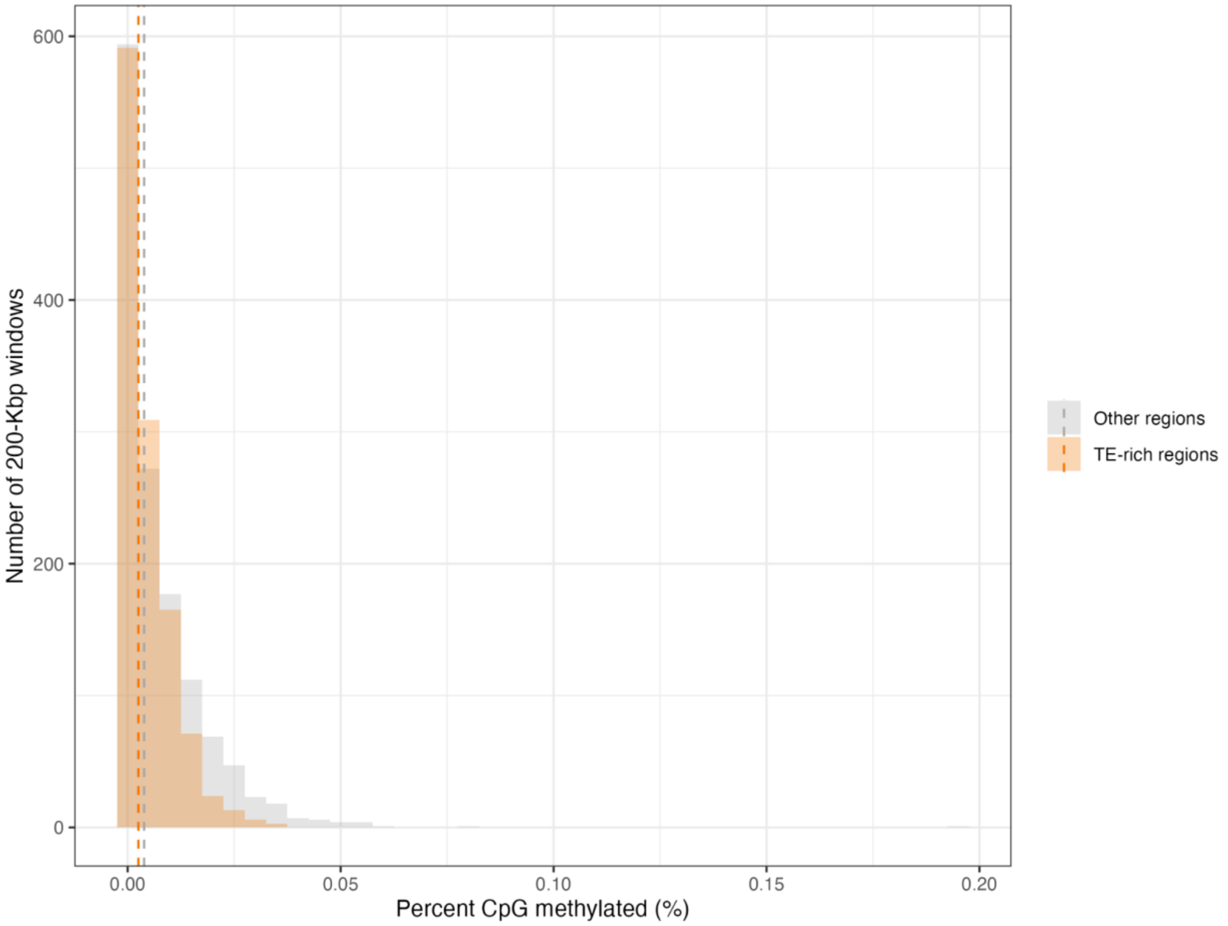


**Supplementary Figure 14: Histogram of percet CpG methylated in 100-Kbp windows in TE-rich regions and other regions.** Dashed lines indicate median value of percent CpG methylated in two regions.

**Supplementary Figure 15**

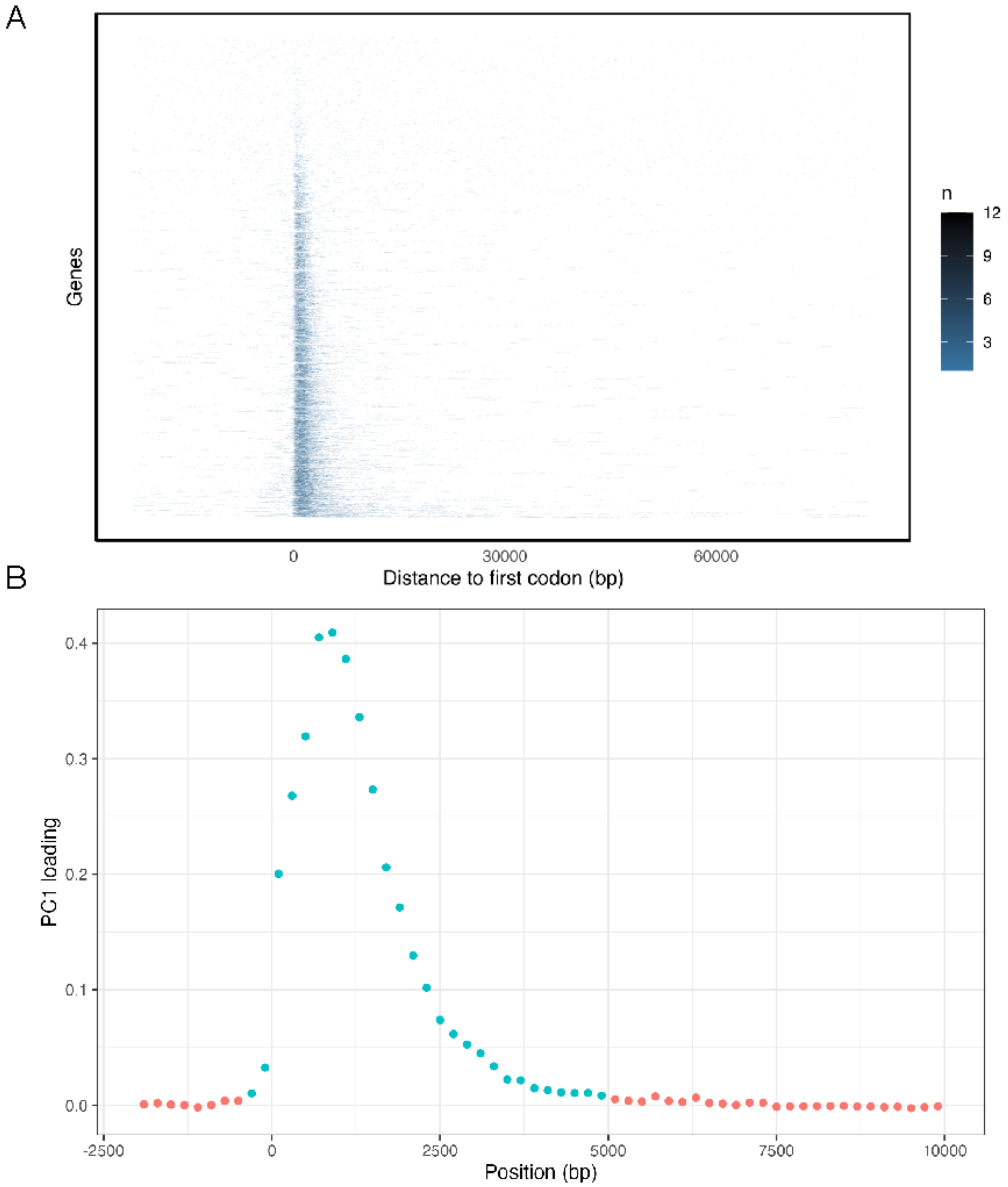


**Supplementary Figure 15: Methylation Pattern around Genes.** (A) Heatmap of the number of methylated CpGs in 500-bp windows. Rows are genes sorted by total number of methylated CpGs near them. (B) PC1 loading of 500-bp windows shows differentially methylated positions near genes in blue (top 5%).

### Supplementary Figure 16

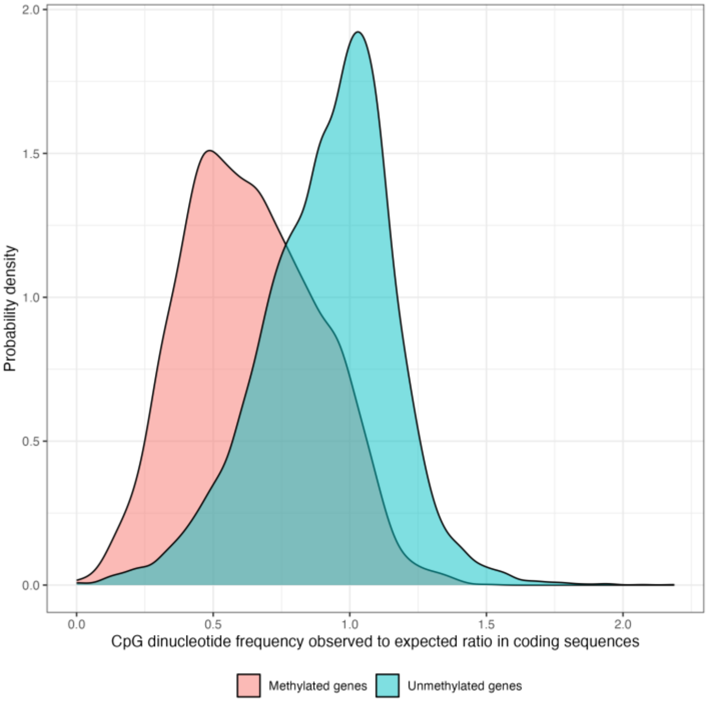


**Supplementary Figure 16: Observed-to-expected CpG ratios for methylated and unmethylated genes.**

#### Supplementary Figure 17

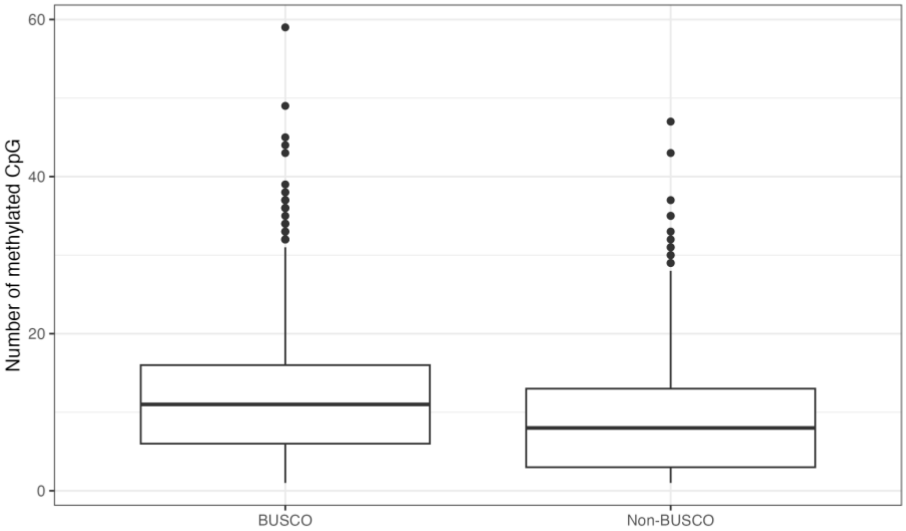


**Supplementary Figure 17: Number of methylated CpGs near BUSCO and non-BUSCO genes.**

#### Supplementary Figure 18

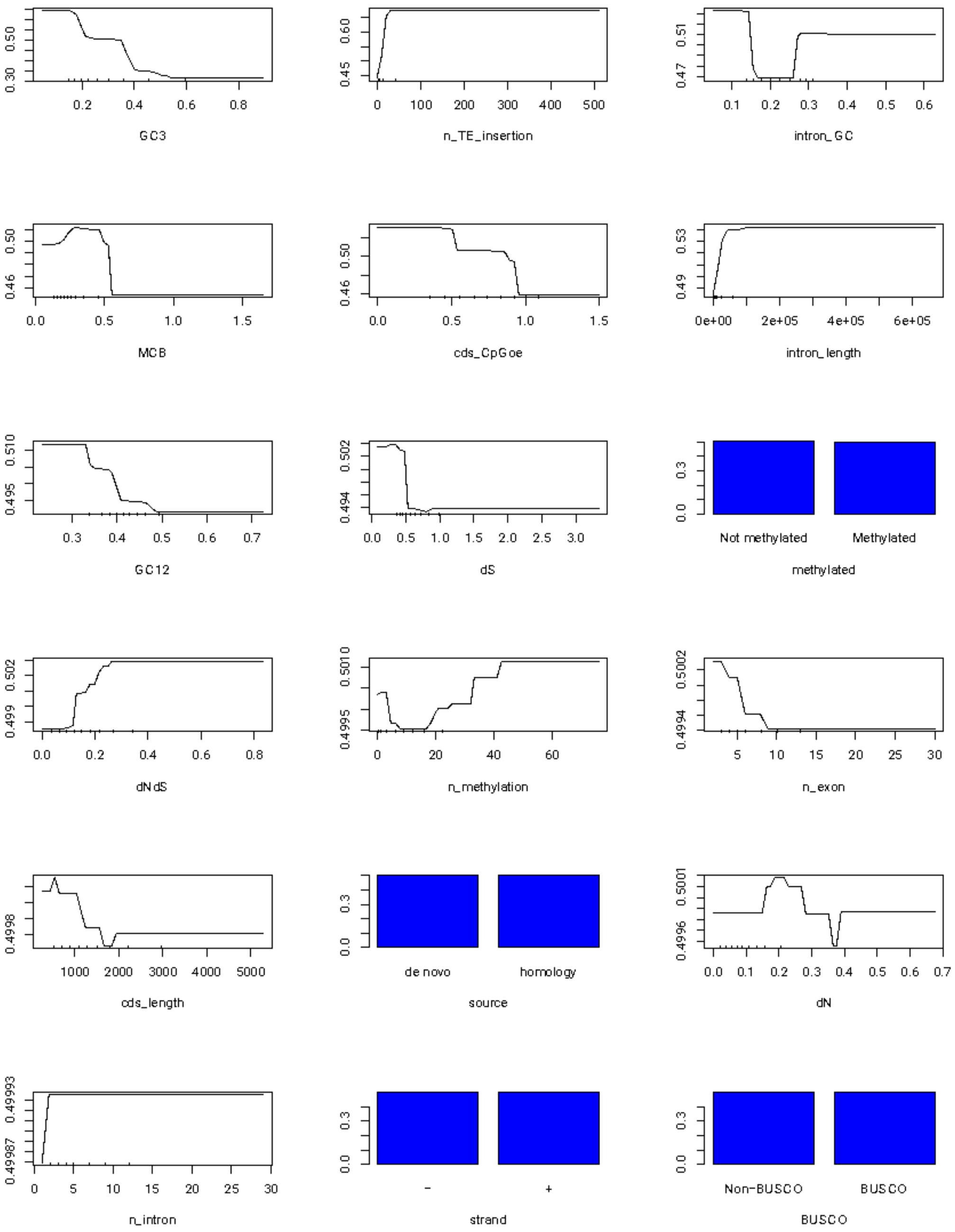


**Supplementary Figure 18: Partial dependence plot of gene features on predicting genes in TE-dense regions.** For numeric variables, hash marks were drawn at the bottom of the plot indicating the deciles. Higher probabilities mean the values of the variable support that the given gene is in TE-rich region.

#### Supplementary Figure 19

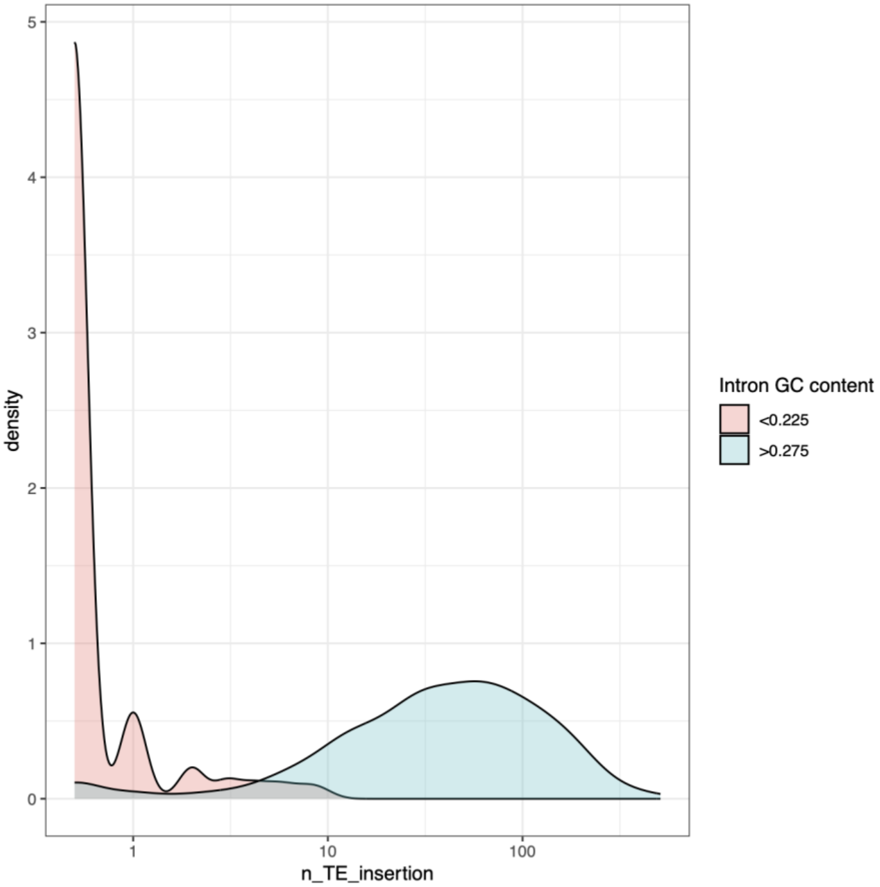


**Supplementary Figure 19: Histogram of the number of TE insertions in genes in TE-rich regions with high and low intron GC content.**

#### Supplementary Figure 20

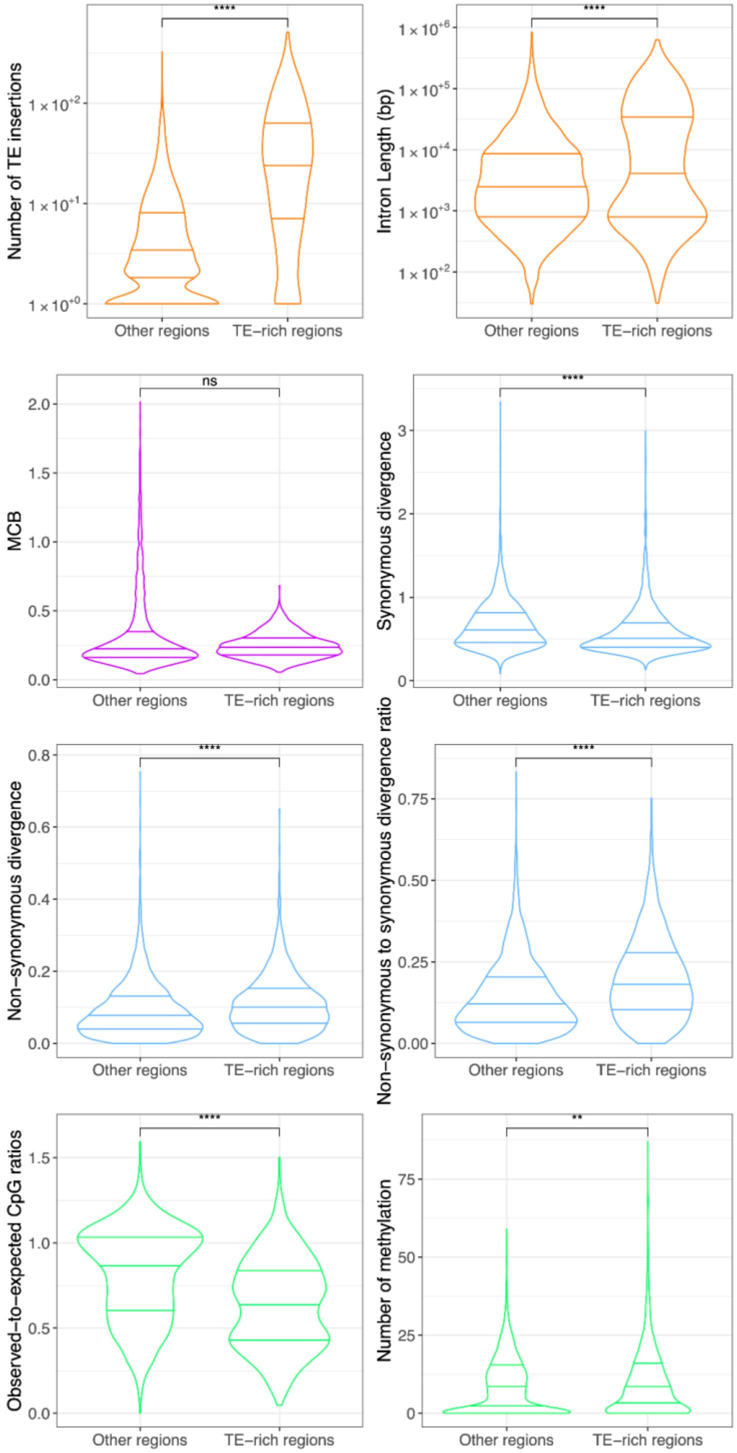


**Supplementary Figure 20: Comparison of other gene features.** Lines indicate 75%, 50% and 25% quantiles from top to bottom.

#### Supplementary Figure 21

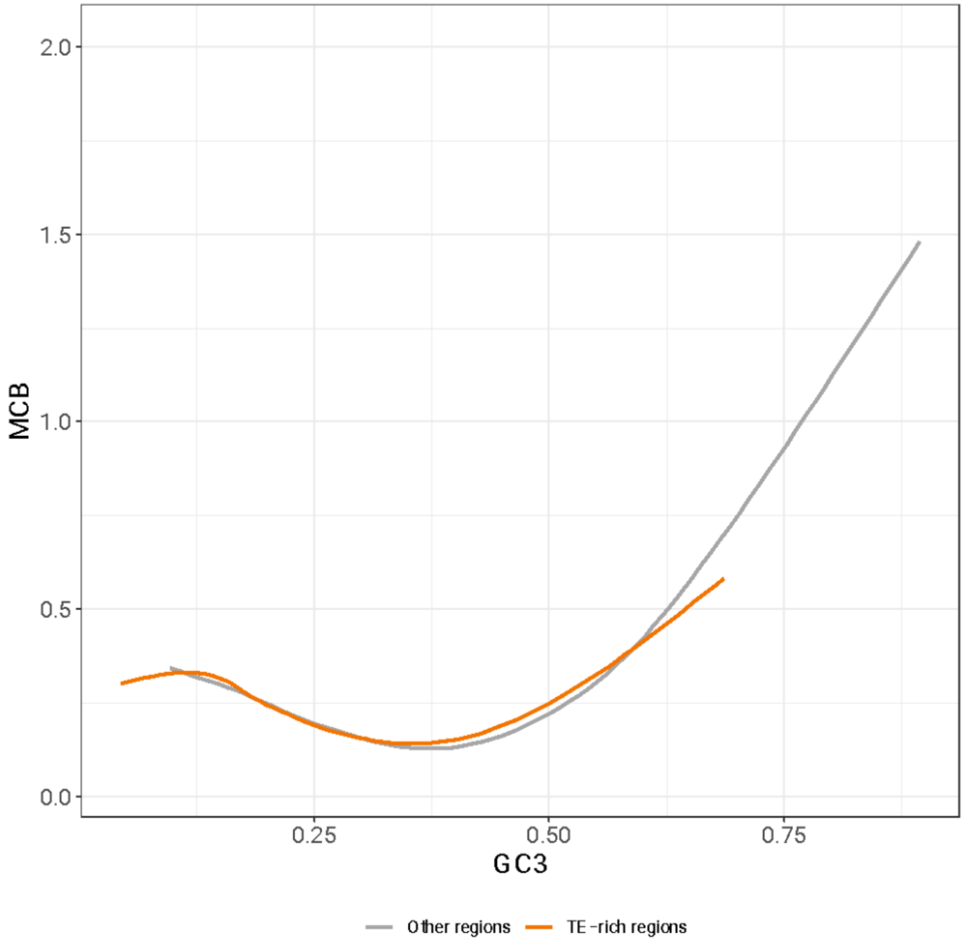


**Supplementary Figure 21: General trend of GC3 versus codon bias.**

**Supplementary Figure 22**

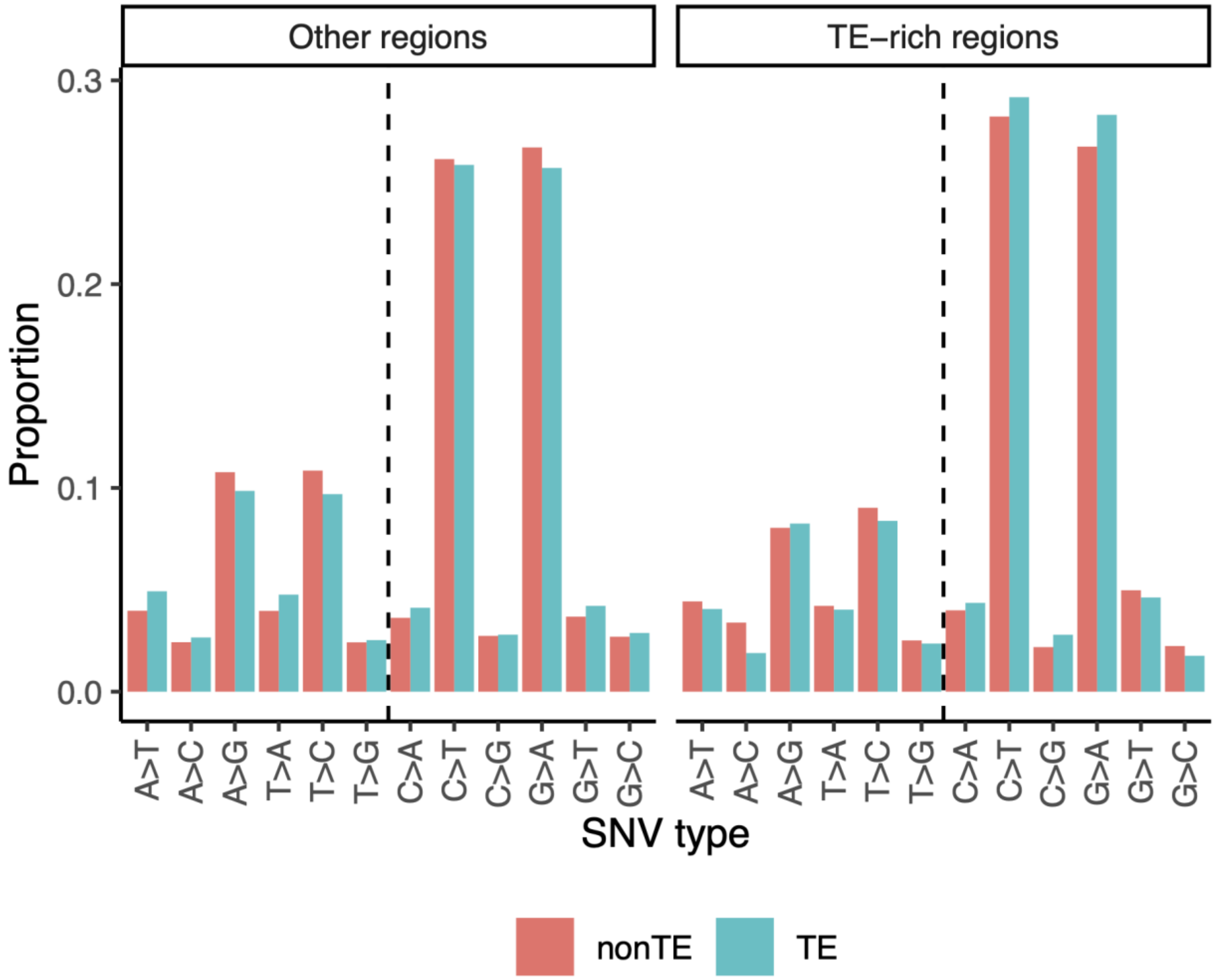


**Supplementary Figure 22: Mutational spectrum of TE and nonTE sequences in TE-rich regions and other regions.** Dashed line indicates two groups of mutations: left to the line are mutations from A and T, and right to the line are mutations from C and G. Color denotes whether the SNV is located in TE sequences.

### Supplementary Tables

#### Supplementary Table 1

| Gene annotation | TE annotation | size (Mbp) |
| --- | --- | --- |
| exon | TE | 0.734771 |
| intron | TE | 54.917487 |
| intergenic | TE | 109.584747 |
| exon | non_TE | 16.818594 |
| intron | non_TE | 115.369164 |
| intergenic | non_TE | 207.31727 |
| Total | | 504.742033 |

**Supplementary Table 1 Genome Annotation Breakdown:** Chromosomal scaffolds are separated into exons, introns and intergenic sequences as well as TE and non-TE sequences.

#### Supplementary Table 2

| Model | Number of unimodal distributions | Number of parameters | Log likelihood | AIC | BIC |
|---|---|---|---|---|---|
| normal | 1 | 2 | 18.63142 | -33.26284 | -30.46072855 |
| exponential | 1 | 1 | 292.3392 | -582.6784 | -581.2773443 |
| normal + normal | 2 | 5 | 11.33955 | -12.6791 | -5.673821371 |
| exponential + exponential | 2 | 3 | 292.3386 | -578.6772 | -574.4740328 |
| exponential + normal | 2 | 4 | 557.8865 | -1107.773 | -1102.168777 |
| beta | 1 | 2 | 560.8332 | -1117.6664 | -1114.864289 |
| beta + beta | 2 | 5 | 1205.041 | -2400.082 | -2393.076721 |
| beta + beta + beta | 3 | 8 | 1205.041 | -2394.082 | -2382.873554 |

**Supplementary Table 2 Model Selection of TE densities.** AIC: Akaike information criterion; BIC: Bayesian information criterion.

#### Supplementary Table 3

| Chromosome | Number of TE-rich windows | Number of TE rich regions | Total size (Mbp) | Proportion of TE-rich regions (%) |
|---|---|---|---|---|
| chr01 | 212 | 3 | 42.4 | 38.57 |
| chr02 | 251 | 4 | 50.2 | 51.02 |
| chr03 | 273 | 2 | 54.6 | 50.73 |
| chr04 | 188 | 1 | 37.6 | 44.6 |
| chr05 | 261 | 2 | 52.2 | 50.27 |

**Supplementary Table 3 Statistics of TE-dense windows and regions per chromosome.**

#### Supplementary Table 4

| Sample | Number of reads before trimming (M) | Number of reads after trimming (M) | Median Mapping Depth | Mapping Ratio (%) |
|---|---|---|---|---|
| W2M.2 | 90.1 | 89.8 | 24 | 99.9 |
| W2M.6 | 96.2 | 95.1 | 24 | 99.5 |
| W2M.8 | 97 | 95.7 | 25 | 99.9 |
| W2M.9 | 86.8 | 85.2 | 22 | 99.5 |
| W2M.12 | 89.4 | 87.4 | 23 | 99.7 |
| W2M.14 | 83.2 | 81.4 | 21 | 99.8 |
| W2M.15 | 89 | 87.2 | 23 | 99.7 |
| W2M.17 | 118.2 | 116 | 30 | 99.8 |

**Supplementary Table 4 Illumina Sequencing and Mapping Statistics.** Median Mapping Depth: median number of reads mapped to a locus. Mapping Ratio: percent of reads mapped.

#### Supplementary Table 5

| Species | Accession | Genome Size (Mb) | Number of protein coding genes |
|---|---|---|---|
| *Anopheles gambiae* | GCF_943734735.2 | 264.5 | 15165 |
| *Apis mellifera* | GCF_003254395.2 | 225.2 | 9935 |
| *Bombyx mori* | GCF_030269925.1 | 461.7 | 13459 |
| *Copidosoma floridanum* | GCF_000648655.2 | 554 | 12143 |
| *Ceratosolen solmsi* | GCF_000503995.1 | 275.78 | 10189 |
| *Drosophila melanogaster* | GCF_000001215.4 | 143.7 | 13962 |
| *Eupristina verticillata* | GWHALOE00000000 | 386.98 | 14281 |
| *Nasonia vitripennis* | GCF_009193385.2 | 297.3 | 13602 |
| *Tribolium castaneum* | GCF_031307605.1 | 241.8 | 12172 |
| *Wiebesia pumilae* | CNA0019285 | 318.19 | 12316 |

**Supplementary Table 5 Reference Genome Annotation Used for GeMoMa.**

#### Supplementary Table 6

| score/aa | Number of genes annotated | Number of complete BUSCO genes | Number of duplicated BUSCO genes | Number of fragmented BUSCO genes | Number of missing BUSCO genes |
|---|---|---|---|---|---|
| 0.75 | 16332 | 5421 | 1331 | 132 | 438 |
| 0.8 | 15984 | 5426 | 1332 | 127 | 438 |
| 0.9 | 15350 | 5435 | 1328 | 122 | 434 |
| 1 | 14812 | 5443 | 1328 | 118 | 430 |
| 1.1 | 14167 | 5446 | 1324 | 118 | 427 |
| 1.2 | 13631 | 5448 | 1315 | 116 | 427 |
| 1.3 | 13140 | 5449 | 1309 | 115 | 427 |
| 1.4 | 12668 | 5453 | 1295 | 113 | 425 |
| 1.5 | 12285 | 5456 | 1291 | 108 | 427 |
| 1.6 | 11916 | 5455 | 1284 | 106 | 430 |
| 1.7 | 11551 | 5454 | 1273 | 106 | 431 |
| 1.8 | 11240 | 5446 | 1258 | 109 | 436 |
| 1.9 | 10936 | 5441 | 1241 | 107 | 443 |
| 2 | 10694 | 5442 | 1230 | 103 | 446 |

**Supplementary Table 6 Parameter Optimization of GeMoMa:** Annotation scores are biased by gene lengths. We tuned the score/aa ratio filtering according to the developers (https://github.com/Jstacs/Jstacs/issues/27) and find 1.5 to be the best parameter value according to the BUSCO completeness.

#### Supplementary Table 7

| CpG | non-BUSCO | BUSCO |
|---|---|---|
| Not methylated | 5153 | 1089 |
| Methylated | 2731 | 3706 |

**Supplementary Table 7**: Patterns of methylation in BUSCO and non-BUSCO genes.

#### Supplementary Table 8

| CpG | intron | exon |
|---|---|---|
| methtlated | 16078 (0.32%) | 44428 (6.32%) |
| not methtlated | 5020968 | 658589 |

Supplementary Table 8: Number of methylated CpGs in introns and exons.
